# MITF maintains genome stability in non-melanocytic cell lineages and suppresses Hippo pathway signaling

**DOI:** 10.1101/2025.06.09.658599

**Authors:** Drifa H. Gudmundsdottir, Adrián López García de Lomana, Thejus B. Venkatesh, Kritika Kirty, Snaevar Sigurdsson, Linda Vidarsdottir, Ramile Dilshat, Erla Sveinbjornsdottir, Snaedis Ragnarsdottir, Daniel H Magnusson, Maria R. Bustos, Eirikur Steingrimsson, Thorkell Gudjonsson, Stefan Sigurdsson

**Affiliations:** Cancer research laboratory, Faculty of medicine, University of Iceland, 102 Reykjavik Iceland; Biomedical center, University of Iceland, 102 Reykjavik Iceland; Department of genetics and molecular medicine University hospital of Iceland, 102 Reykjavik Iceland; Biotech Research & Innovation Center, University of Copenhagen, 2200 Copenhagen Denmark

## Abstract

Microphthalmia-associated transcription factor (MITF) is crucial for development and survival of melanocytes and serves as a lineage-specific oncogene that is amplified in 10-20% of melanomas. The role of MITF in pathways maintaining genome integrity, such as DNA replication, DNA repair and mitosis has been extensively studied in melanocytes. In addition to its pro-survival role in melanoma, recent studies have shown that MITF expression has important implications for cancer progression and treatment in other cancer types. Nevertheless, studies on the role of MITF in other tissues are scarce. Here, we show that depletion of MITF causes genome instability in non-melanocytic cell lineages, which results in activation of P53, cell cycle arrest and apoptosis. Moreover, we show that P53 activation in MITF depleted cells is dependent on LATS2, a key kinase in the Hippo pathway. Finally, we show that this LATS2 mediated upregulation of P53 is ATR dependent. Collectively, this study highlights the role of MITF as a genome maintenance factor beyond the melanocyte lineage, which might contribute to the tumor suppressive function of MITF.

## Introduction

MITF has an established role as a master regulator of melanocytes and a driver of melanoma progression(1–3). Importantly, MITF controls melanoma plasticity, by switching between proliferating and metastatic behavior of melanoma cells(4, 5) Increased MITF expression has been associated with poor prognosis in melanoma patients(6), underlining the clinical relevance of MITF. The MITF gene encodes nine isoforms, which differ in tissue specificity and transcriptional targets(3, 7). Although the melanocyte specific M-isoform has been extensively studied, characterization of the other isoforms is lacking.

In addition to its oncogenic function in melanoma, deregulation of MITF expression has important implications in various cancer types(8, 9) highlighting the clinical relevance of MITF in non-melanocytic cell linages. MITF expression significantly impacts survival and resistance to CDK4/6 inhibitors in breast cancer(10) In non-small cell lung cancer, high MITF expression correlates with improved prognosis. In line with that, silencing MITF in mouse models promotes metastasis and tumorigenesis(9), suggesting MITF suppresses tumour formation in lung cancer. In contrast to the oncogenic role of MITF, the mechanism underlying the tumour suppressive function of MITF remains unclear.

Genome instability refers to high frequency of genetic alterations and is one of the major hallmarks of cancer. To maintain genome stability, dividing cells must faithfully duplicate their genome in each cell cycle and ensure an equal distribution of chromosomes to daughter cells. Replication stress is a broad term, which covers a variety of stresses that interfere with DNA replication and is recognized as the major source of genome instability in cancer(11, 12). Replication stress can cause the DNA replication machinery to halt, which increases the risk of replication fork collapse and DNA double strand break (DSB) formation(13, 14). Failure to complete DNA replication in S-phase increases the risk of cells entering mitosis with under-replicated DNA (urDNA), where it can cause aberrant segregation of chromosomes and DNA breakage(15). Previous studies have shown that MITF plays a role in maintaining genome stability by controlling DNA replication, DNA repair and mitosis in the melanocyte lineage(16–18).

The Hippo pathway is known to play a role in tissue growth through regulation of cell proliferation and apoptosis(19, 20). Since its discovery, it has been shown that the Hippo pathway mediates cell cycle arrest and cell death in response to various stresses, including replication- and mitotic stress(21–23). Therefore, it is not surprising that aberrant regulation of Hippo pathway signaling has been associated with tumorigenesis and tumor progression(24–27). *Large tumor suppressor kinase 2* (LATS2) plays a key role in Hippo pathway signaling in response to replication stress and promotes P53 DNA binding which results in transcription of apoptosis-promoting genes(21, 28, 29). Further, in response to stress, LATS2 has been shown to promote rapid P53 protein accumulation, by translocating to the nucleus where it binds and inhibits *Mouse double minute 2 homolog* (MDM2), an E3 ubiquitin ligase responsible for P53 proteasomal degradation(21, 23, 28, 29).

In this study, we show that depletion of MITF in non-melanocyte cell lines negatively affects cell proliferation and DNA replication, leading to genome instability. We also show P53 activation upon MITF knockdown, which results in cell cycle arrest and apoptosis. Furthermore, we show that the observed P53 activation is mediated through LATS2, thereby uncovering a previously unrecognized connection between MITF and members of the Hippo pathway. Overall, our findings suggest that MITF plays an important role in maintaining genome stability in non-melanocytic cell linages, which might be important for the tumor suppressive function of MITF.

## Materials and methods

### Reagents

Aphidicolin (Aph) (Abcam, ab142400). KaryoMAX^TM^ Colcemid^TM^ (Thermo, 15212012). ATM inhibitor KU55933 (Sigma Aldrich, 587871-26-9). ATR inhibitor VE-821 (Sigma Aldrich, 1232410-49-9), AuroraA inhibitor JB300 (Tocris Bioscience, 7837/5).

Nocodazole (Sigma Aldrich, SML1665). Doxycycline (Sigma Aldrich, D9891-1G).

Doxorubicin (abcam, ab120629). Etoposide (Sigma Aldrich, E1383). Karymax^TM^ colcemid (Thermo, 15212012).

Following antibodies were used in this study: anti-MITF C5 (abcam, ab12039), anti-53BP1 (Santa cruz, sc-22760), anti-γtubulin (abcam, ab11316), anti-p53 (Cell Signaling, 2527), anti- SMC1 (abcam, ab9262), anti-γH2AX (abcam, ab22551), anti-MYCtag (Cell Signaling, 2272), anti-CyclinA (Santa Cruz, sc-271682), HRP 2° antibodies (Santa cruz) (anti-mouse (sc- 2096), anti-rabbit (sc-2313)), Alexa Fluor 488 anti-mouse 2° antibody (Life technologies, A21121), Alexa Fluor 488 anti-rabbit 2° antibody (Invitrogen, A32731), Alexa Fluor 555 anti-rabbit 2° antibody (Invitrogen, A21428), Alexa Fluor 555 anti-mouse 2° antibody (Invitrogen, A21422), Alexa Fluor 647 anti-mouse 2° antibody (Invitrogen, A21235).

Following plasmid was used in this study: pClneoMYC-LATS2 (66852) (Addgene).

Kits and other reagents: Lipofectamine^®^ RNAiMAX (Thermo Fisher, 13778075), Lipofectamine^®^ LTX with PLUS^TM^ reagent (Thermo Fisher, 15338100), Pierce^TM^ 16% Formaldehyde (W/V), methanol free (Thermo Fisher, 28906), Click-It^TM^ EdU kit (Thermo, C10419), RNeasy Mini kit (Qiagen, 74104), TRI reagent (Thermo, AM9738), Superscript II (Thermo, 18064022), SYBR green^®^ (Thermo, 25742), CellTrace^TM^ Violet cell Proliferation kit (Invitrogen, C34571), caspase-3/7 red dye (Sartorius, 4704).

Specialized commercial instruments: Olympus FV 1200 Confocal microscope, Attune NxT Flow cytometer (Thermo Fisher), Navios Ex (Beckman Coulter), CCD camera (BIO-RAD ChemiDoc^TM^ XRS+), Incucyte^®^ S3 live-cell imager, CFX Real-time PCR Detection system (Bio-Rad).

### Biological resources

The following cell lines were used in this study: U2OS (human osteosarcoma), 624mel (human melanoma), Sk-Mel-28 (human melanoma), HeLa (human cervical cancer), A549 (human lung adenocarcinoma), D492 (human breast epithelia), SW-1353 (chondrosarcoma).

### Cell culture

Sk-Mel-28, 624mel and derived cell lines were cultured in RPMI 1640 medium and U2OS, HeLa, SW1353 and A549 cells were cultured in Dulbebecco’s modified Eagle medium (DMEM (1x) + Glutamax^TM^ -I medium (Thermo, 31966047)). All cancer cells were cultured with 10% fetal bovine serum (FBS) (Thermo, A5256801) and antibiotics (Penicillin 2 units/500mL and Streptomycin 2μg/500mL (Thermo, 15070-063)). The non-cancer cell line D492 was cultured in a H14 medium (DMEM/F-12 media (Thermo, 31330038), 250 ng/ml insulin (Sigma, 16634), 10 µg/ml transferrin (Sigma, T8158), 10 ng/ml EGF (PeproTech, 100- 15), 2.6 ng/ml sodium selenite (Sigma, S5261), 0.1 nM estradiol (Sigma, E2758), 0.5 µg/ml hydrocortisone (Sigma, H0888) and 5 µg/ml prolactin (PeproTech, 100-07). Cells were cultured in a humidified incubator at 37°C with 5% CO2. Cells were Mycoplasma tested on regular basis.

### siRNA and plasmid transfection

Lipofectamine^®^ RNAiMAX (Thermo Fisher) was used for siRNA transfection according to manufactureŕs recommendations. Briefly, transfection reagent was removed 24 h after transfection and fresh media added. The following siRNAs (Thermo Fisher, Silencer^®^ and Silencer select^TM^) were used in this study: siMITF (s8791 (#1), s8792(#2)), siP53 (s606), siLATS2 (s25503), siLATS1 (s17393), siBRCA1(s458), siBRCA2 (s2085), si53BP1 (107785), siBLM (s1997) and siCON No.1.

Lipofectamine^®^ LTX with PLUS^TM^ reagent (Thermo Fisher) was used for plasmid transfections according to manufactureŕs recommendations.

### Fluorescent microscope imaging

For immunofluorescence (IF) staining, cells were grown on autoclaved glass cover slips. Cells were fixed using 4% formaldehyde (diluted Pierce^TM^ 16% Formaldehyde (W/V), methanol free (Thermo Fisher, 28906)), for 15 min at RT. Cells were permeabilized in 0.2% Triton-X (Merck, 108643) in PBS for 4 min at RTe and blocked in blocking buffer (DMEM (1x) + Glutamax^TM^ -I medium + 10%FBS) for 30 min. Samples were incubated in primary antibody diluted in blocking buffer for 90 min at RT. Secondary antibody and 4’, 6-diamidino-2- phenylindole (DAPI)(Sigma, D9542)(1μg/mL) staining was performed at RT for 60 min. Samples were washed in distilled water following two PBS washes and then dried. Coverslips were placed on a drop (4µL) of mounting medium (Santa Cruz, sc516212) on a microscope slide. Click-It^TM^ EdU kit (Thermo, C10419) was used to mark cells that were actively replicating DNA. Olympus FV 1200 Confocal microscope was used for imaging. Automated image analysis was performed using the CellProfiler^TM^ software. Protein intensity was measured using the mean intensity output of the MeasureObjectIntensity module. Nuclear and foci quantification was performed using the IdentifyPrimaryObject module. For image acquisition the 405 nm laser was used to identify DAPI stained nuclei, for visualization of the targets fluorescent secondary antibodies ere used (Alexa Fluor 488/555/647) in combination with image acquisition with lasers emitting light at 473 nm, 543 nm and 635nm.

### Flow cytometry

DNA content was stained with 7-aminoactinomycin D (7AAD)(Thermo, A1310) or DAPI for flow cytometry analyses. Before staining cells were fixed using 70% ethanol. 1x10^6^ cells were resuspended in 500µL of ice-cold PBS, 1.2mL ice cold ethanol was added drop wise while vortexing, samples were then incubated for at least 3 h at -20°C to complete fixation. After fixation samples were washed in PBS and resuspended in 300µL of FACS staining buffer (0.1% Triton-X in 1xPBS) with 7AAD (25µg/mL(for 5 x 10^5^ cells)) or DAPI (5μg/mL (for 5 x 10^5^ cells)). Samples were incubated in the dark for 30 min at RT. Samples were analyzed using the Attune NxT Flow cytometer (Thermo Fisher), Navios Ex (Beckman Coulter), and Flowjo^TM^ software. Dye intensity was measured using Attune Flow cytometer (for DAPI: Ex. Laser: 405, Em. Filter: 440/50. for 7AAD: Ex. Laser: 561, Em. Filter: 620/15) (Thermo Fisher) (Ex. laser:405, Em. filter:440/50) and Flowjo (Thermo Fisher) software.

### RNA sequencing

RNA was isolated using TRI reagent (Thermo, AM9738). The RNA was DNase1 treated (Qiagen) and purified using RNeasy Mini kit (Qiagen). The RNA samples were analyzed using Bioanalyzer (Agilent Technologies), all samples were found to have RNA integrity of 9.8 and higher. Paired and library sequencing was carried out on the Illumina platform. For gene expression quantification, we first cleaned original FASTQ files using Trimmomatic version 0.39 using default parameters [doi:10.1093/bioinformatics/btu170](30). Next, we quantified gene expression from generated FASTQ files using kallisto version 0.50.1 [doi:10.1038/nbt.3519] and the Ensembl Homo sapiens reference transcriptome (version 113). For differential gene expression analysis, first, we filtered out genes that were expressed at very low levels (max expression < 1 TPM) and less than 20 reads in expression difference across conditions. Then, we used DESeq2 version 1.26.0 [doi:10.1186/s13059- 014-0550-8] to determine statistically significant differentially expressed genes (DEGs) across experimental conditions (Benjamini–Hochberg correction α = 0.05; adjusted P < 0.05; |log2 FC| < 1). For functional enrichment analysis, we used clusterProfiler [doi:10.1038/s41596-024-01020-z] on Reactome Pathway ontology [doi:10.1093/nar/gkad1025] to functionally characterize DEG sets. We accepted as enriched terms those with a *P* < 0.05 after multiple test correction. Code to reproduce all gene expression quantitative analysis is publicly available within GitHub repository https://github.com/adelomana/BMCBF/tree/main/research/025_isafjordur

### Metaphase spreads

Cells were cultured on 6-well plates and harvested seven days post siRNA treatment. To collect cells in metaphase, Colcemid was added to each sample 4 h before harvest. Pelleted cells were dissolved in a small volume of media, 2mL hypotonic solution (0.075M KCl) (37°C) was then added dropwise to each sample while gently vortexing. Samples were incubated at 37°C for 10 min. Cells were pelleted and dissolved in small volumes of the supernatant. 3mL of cold fixative (3:1 methanol and acetic acid) was added dropwise to each sample while vortexing, samples were then incubated on ice for 10 min. The fixative steps were repeated 2-3 times until the supernatant became clear. The pellets were resuspended in a small volume of fixative.

### Western blotting

Whole cell lysate from cell culture (grown on 6 well plates) were prepared by adding 100µL of 2X Laemmli Sample Buffer (Santa Cruz, sc286963) to each sample and the cells scraped and moved to an Eppendorf tube. Benzonase (1 ng/mL) (Sigma, E1015-5KU) was added before samples were heated at 95°C 7-8 min. Samples were blotted onto Nitrocellulose membrane (0.4µM) (Santa Cruz, sc3724). Membranes were incubated in primary antibodies O/N at 4° in PBS with 5% milk. Secondary antibodies were incubated for 60 min at RT in PBS with 5% milk. Membranes were treated with luminol reagent (Santa Cruz, sc2048) for 60 sec, and images acquired using CCD camera (BIO-RAD ChemiDoc^TM^ XRS+).

### qPCR

RNA was isolated using TRI reagent (Thermo, AM9738) and NanoDrop one (Thermo Scientific) was used to assess RNA concentration and purity. RNA samples were reverse transcribed using Superscript II (Thermo, 18064022) according to the manufacturer’s protocol, using 1ng isolated RNA and oligo (dT) in a total reaction volume of 19 μL, followed by RNase treatment (NEB, M0314L). SYBR green^®^ (Thermo, 25742) amplification detection was used for the qPCR and β-actin was used to normalize data. All experiments were performed in technical triplicates and at least 3 biological replicates. Primer sequences available in supplementary data S2. CFX Real-time PCR Detection system (Bio-Rad) was used for qPCR experiments. For fold change calculations the ΔΔCT method was used.

### Cell proliferation assays

To assess cell proliferation CellTrace^TM^ Violet cell Proliferation kit (Invitrogen, C34571) was used. The manufacturer protocol was used for staining cells. Dye intensity was compared between treatments, where higher intensity indicated fewer cell divisions. Dye intensity was measured using Attune Flow Cytometer (Thermo Fisher) (Ex. laser:405, Em. filter:440/50) and FlowJo (Thermo Fisher) software.

### Apoptosis assays

To analyze apoptosis, we used Incucyte^®^ S3 live-cell imager (Sartorius) and caspase-3/7 red dye (Sartorius, 4704). Cells were reverse transfected in 96 well plates (3000 cells/well). After 24 h siRNA treatment the caspase-3/7 dye (0.5 μM) was added. Cell confluency was obtained using the Incucyte^®^ S3 software and Caspase-3/7 positive cells were counted manually.

### Survival assays

For survival and drug sensitivity assays we used the Incucyte^®^ S3 live-cell imager where cell confluency was analyzed using the Incucyte^®^ S3 software after 168h siRNA treatment.

### Invasion assay

8μm pore transwell filters were used for the invasion assays (Corning). 24h after siRNA transfection, 40.000 cells were seeded on Matrigel (3% in serum free media)(Corning, 354230). Cells were given 20h to invade through the Matrigel towards media containing 10% FBS. Invading cells were fixed in 4% formaldehyde for 5 min at RT and permeabilized in 0.1% Triton-X for 5 min at 4°C. Cells were stained in DAPI (1μg/mL) diluted in PBS for 30 min at RT in the dark. Images were acquired using EVOS FL fluorescent microscope (Thermo Fisher Scientific).

### Statistical analyses

To compare the means of two groups, unpaired two-tailed Student’s T-test was used to determine statistical significance. To compare the means of more than two groups, one-way ANOVA was used. Error bars represented as the standard deviation. At least 3 biological replicates were used for each statistical analysis. Throughout the paper, statistical significance was presented as follows: \**P* <0.05, \*\**P* <0.01, \*\*\**P* <0.001, \*\*\**P* <0.0001.

### Data availability/novel programs, software, algorithms

Softwares and programs used in this study: CellProfiler cell image analysis software (https://cellprofiler.org/), FlowJo (v10) flow cytometry analysis software, Kaluza analysis 2.2 flow cytometry analysis software, Fiji image processing software (https://fiji.sc/).

### Web/site/database referencing

Following website was used in this manuscript: The Human Protein Atlas (HPA) (https://www.proteinatlas.org; https://www.proteinatlas.org/ENSG00000187098MITF/subcellular).

## Results

### MITF promotes genome stability in non-melanocyte cell lines

The role of the melanocyte specific MITF-M isoform in melanocyte development and melanoma has been extensively studied. However, several other MITF isoforms have been found to be ubiquitously expressed in various tissue types, although their functional role remains to be characterized. Here, we hypothesized that the MITF’s role, as a genome maintenance factor is not restricted to the melanocyte lineage. Firstly, we took advantage of The Human Protein Atlas (HPA) in order to determine the mRNA expression of MITF across tissue types. According to the database, MITF is expressed in cell lines originating from several different tissues (Figure 1A). For subsequent experiments, we selected four non-melanocyte cell lines (HeLa (cervical cancer), U2OS (osteosarcoma), SW-1353 (chondrosarcoma) and A549 (lung adenocarcinoma)), which according to HPA have MITF expression greater than 10 nTPM (normalized transcripts per million). To confirm specific MITF mRNA expression in the selected cell lines we performed siRNA mediated MITF knockdowns. The results confirmed MITF mRNA expression in all cell lines tested. (Figure 1B). Abnormal gene expression is commonly seen in cancer cell lines and therefore MITF expression was also measured in the non-cancer breast epithelia cell line, D492(31) (Figure 1B). Next, we compared the MITF mRNA expression values in the four cancer cell lines and the non-cancer line to the melanoma cell line (624mel). As expected, and in agreement with the HPA data, this revealed low, but detectable expression of MITF in the non-melanocyte cell lines, compared to the melanoma cell line (Figure 1C). Collectively this data confirms that MITF is expressed in the non-melanocyte cell lines tested, in low levels compared to the melanoma cell line 624mel.

**Figure 1.**
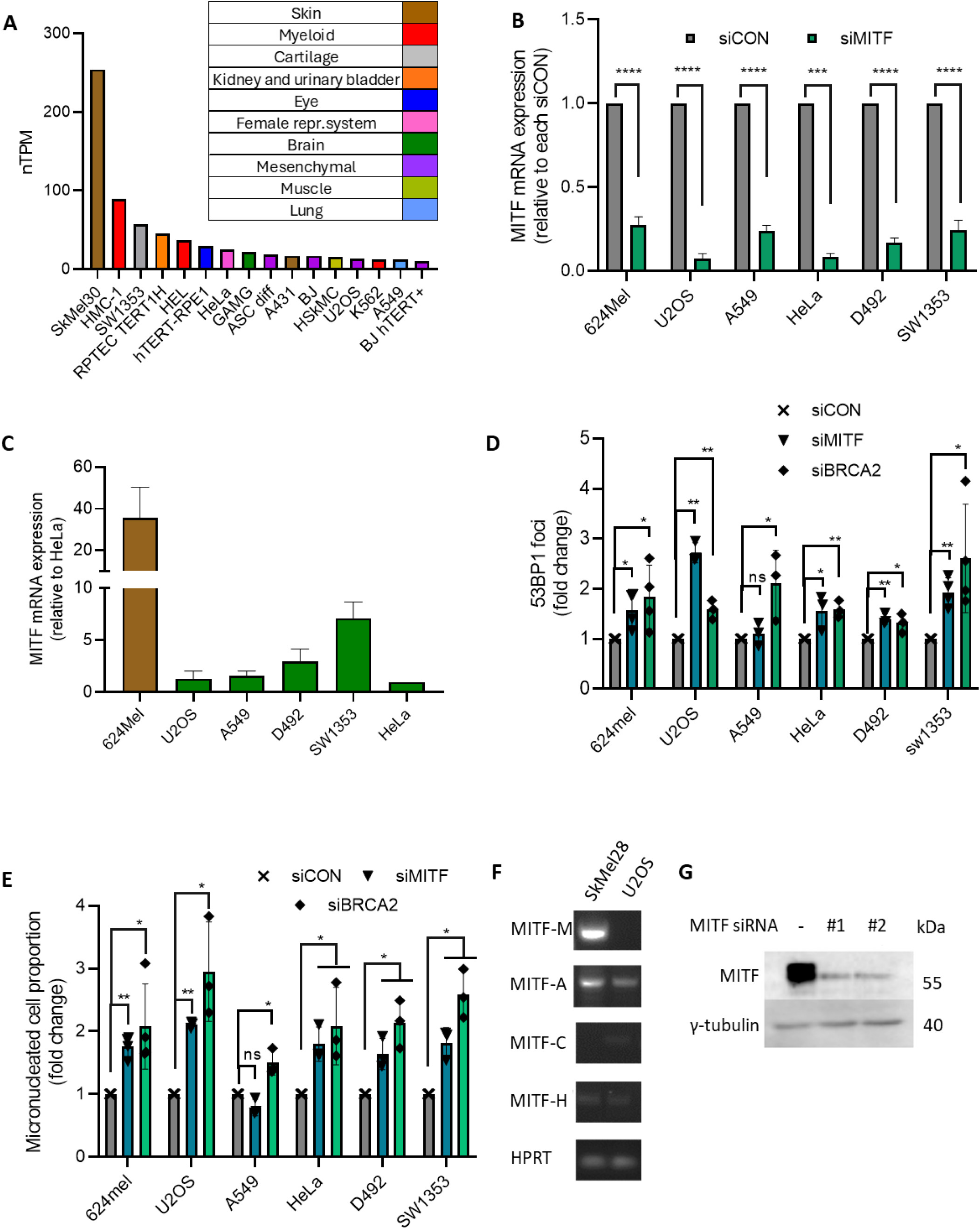
MITF knockdown causes genome instability in non-melanocyte cell lines. **a** MITF normalized transcripts per million (nTPM) in cell lines originated from the indicated tissue types, data obtained from The Human Protein Atlas. **b** MITF mRNA expression confirmed in RNA samples extracted from four non-melanocyte cancer cell lines (U2OS, A549, HeLa and SW1353), a non-cancer breast epithelia cell line (D492) and one melanoma cell line (624mel). Cells were treated with the indicated siRNAs for 48h followed by real time qPCR analysis (unpaired t-test, n=3). **c** Comparison of mRNA expression levels in a melanoma cell line and non-melanocyte cell lines using real-time qPCR analysis. Expression values are shown as fold change, relative to HeLa expression values (n=2). **d** Quantification of nuclear 53BP1 foci in the same cell lines as in b, following treatment with the indicated siRNAs for 72h and immunostaining with 53BP1 specific antibody. The bar chart shows the fold change in 53BP1 foci formation compared to control cells for all five cell lines (unpaired t-test, n=3- 4). **e** Analysis of the proportion of cells with micronuclei in the same cell lines as in b, following 72h treatment with the indicated siRNAs and DAPI staining to visualize nuclear cells. The bar plot shows fold change in the proportion of cells with micronuclei compared to control cells for each cell line (unpaired t-test, n=3-4). **f** Semi-quantitative PCR bands obtained from MITF isoform specific primers. **g** MITF protein expression in U2OS cells treated with control and two independent MITF siRNAs, analyzed by western blot of whole cell extracts using the indicated antibodies. Data presented as mean ± SD. \**P* <0.05, \*\**P* <0.01, \*\*\**P* <0.001, \*\*\*\**P* <0.0001.

In order to determine the impact of MITF expression on genome stability in melanoma and non-melanoma cell lines, the number of 53BP1 foci (marker for DNA DSBs)(32, 33) and the proportion of cells with micronuclei (marker for chromosome instability)(34, 35) after MITF knockdown was analyzed. MITF depletion resulted in a significant increase in 53BP1 foci and micronucleated cell counts in all cell lines tested with the exception of A549 (Figure 1B, D, E and Supplementary Figures S1A-D). This strongly suggests that MITF plays a role in maintaining genome stability, which extends beyond the melanocyte lineage.

### MITF in osteosarcoma cells supports DNA repair and cell cycle regulation

Considering the strong indications of cellular stress detected in MITF depleted U2OS cells, we were prompted to further estimate the degree of genome instability in this particular cell line. We confirmed MITF protein expression in U2OS cells using western blot analysis (Figure 1G). Isoform expression analysis revealed that the MITF-A isoform, which is the longest MITF isoform, is predominantly expressed in U2OS, although low MITF-H expression was also detected. As expected, the melanocyte specific M isoform was not detected in U2OS cells Figure 1F and Supplementary Data S1).

To evaluate the impact of MITF on cell integrity, we performed metaphase spreads to measure the level of chromosomal instability in MITF depleted cells. The metaphase spreads revealed that MITF knockdown resulted in a significant increase in chromosome aberrations compared to control cells, which was a similar increase as was observed with BRCA2 knockdown or induction of mild replication stress by low dose DNA polymerase inhibitor treatment (aphidicolin 200 nM for 24h) (Figure 2A and Supplementary Figure S2A).

**Figure 2.**
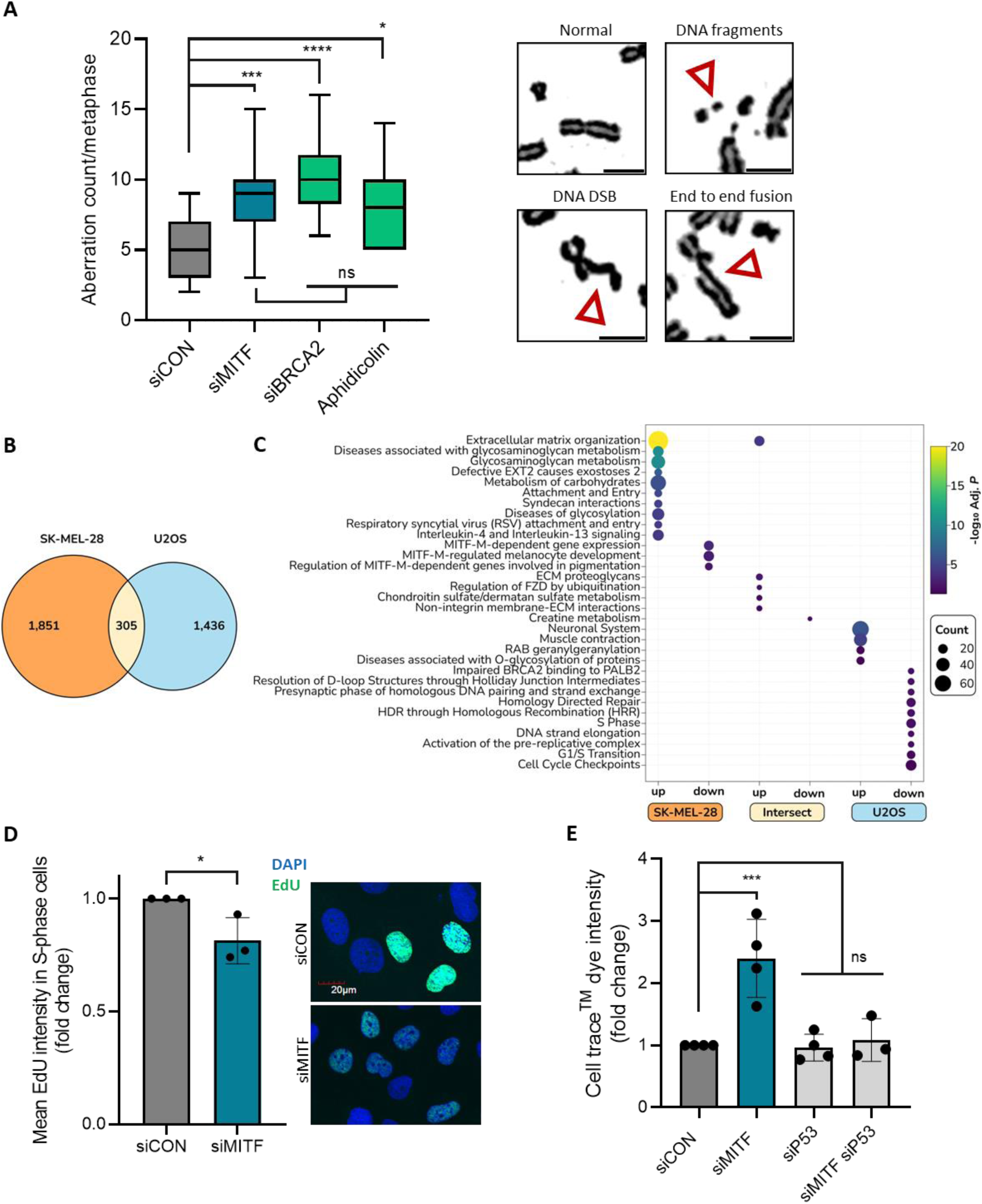
MITF knockdown has an impact on genome stability, DNA replication and cell proliferation in U2OS cells. **a** Quantification of chromosome aberrations (right panel shows examples of aberrations observed) in metaphase spreads samples after seven-day siRNA (siCON, siMITF and siBRCA2) treatment or 24h Aphidicolin treatment (200 nM). To enrich for cells in metaphase, samples were treated with the mitotic inhibitor Colcemid four hours prior to fixation. Scale: 10μM (Welch’s test, n=7-23). **b** Venn diagram comparing differentially expressed gene sets responding to MITF loss in two different cell lines. **c** Dot plot showing a non-redundant summary of functionally enriched pathways for the different gene sets indicated. **d** Mean EdU intensity in S-phase cells after 48h siRNA treatment (siCON and siMITF) and a 30-minute incubation with EdU, followed by a click chemistry staining of EdU and DAPI staining to visualize nuclear cells. Scale: 20μM (unpaired t-test, n=3). **e** Violet blue dye intensity after five-day siRNA treatment (siCON, siMITF, siP53 and siMITF + siP53). Prior to siRNA transfection, cells were stained with a Violet blue cell trace^TM^ dye. Dye intensity was obtained using flow cytometry, higher intensity indicating fewer cell cycles (one-way ANOVA, n=3). Data presented as mean ± SD. \**P* <0.05, \*\**P* <0.01, \*\*\**P* <0.001, \*\*\*\**P* <0.0001.

Therefore, we sought to characterize the molecular mechanisms behind the observed cellular response to MITF loss in U2OS cells. Towards this goal, we compared the transcriptional response to MITF loss in U2OS with respect to its loss in a melanocytic background cell line, SK-MEL-28 specifically. We identified (see Methods for details) a similar number of differentially expressed genes (DEGs) in both cell lines upon MITF loss: 2, 156 DEGs in SK-MEL-28 and 1, 741 DEGs in U2OS (Supplementary table S1, Supplementary Figure S2B). Yet the intersection between these two sets is relatively small: upon perturbation only 305 DEGs (14% of DEGs in SK-MEL-28) are shared between the two cell lines (Fig. 2B). Thus, we hypothesized that the molecular mechanisms behind the two transcriptional responses may be different. To further characterize the perturbed functions in each cell context, we performed functional analysis on the DEGs particular and common to each cell line (Supplementary table S2). We identified two clearly distinct responses (Fig. 2C). The response in SK-MEL-28 is characterized by the upregulation of genes involved in the remodeling of the extracellular matrix as supported by pathway terms such as *Extracellular matrix organization* (*P* < 2 × 10^-21^), *Metabolism of carbohydrates* (*P* < 2 × 10^-6^) and *Syndecan interactions* (*P* < 2 × 10^-5^). The set of downregulated genes in SK-MEL-28 is enriched on MITF-specific targets only such as: *MITF-M-dependent gene expression* (*P* < 5 × 10^-4^), *MITF-M-regulated melanocyte development* (*P* < 2 × 10^-3^) and *Regulation of MITF-M-dependent genes involved in pigmentation* (*P* < 5 × 10^-5^).

However, the transcriptional response in U2OS is unequivocally different. Most of the significantly enriched pathways in the downregulated DEG set (18 out of a total of 25 enriched pathways) elicits to DNA repair functions such as *Impaired BRCA2 binding to PALB2* (*P* < 7 × 10^-3^), *Homology Directed Repair* (*P* < 8 × 10^-3^) and *Diseases of DNA repair* (*P* < 2 × 10^-2^). Six other significantly enriched pathways revolve around cell cycle progression, as supported by the terms *S Phase* (*P* < 8 × 10^-3^), *G1/S Transition* (*P* < 4 × 10^-2^) and *Cell Cycle Checkpoints* (*P* < 4 × 10^-2^) with key genes such as *MCM4, ORC1, FEN1, RFC1, ORC*1 and *CDT1* as clear hallmarks of replication. Altogether, these results indicate that the response to MITF loss in U2OS is dissimilar compared to a melanocytic cell background, and it is indeed markedly characterized in U2OS by the downregulation of genes involved in DNA repair and replication.

Next, we aimed to experimentally validate the impact of MITF on DNA repair and replication in the U2OS cell line. To determine the proportion of DNA replicating cells, U2OS cells were treated with a short pulse of the thymidine analogue EdU, followed by image analysis to determine the rate of EdU incorporation. Interestingly, we detected a drop in EdU incorporation, specifically, a lower mean intensity of Edu in the S-phase of MITF depleted cells compared to control cells, indicative of reduced DNA replication (Figure 2D). Additionally, it is well known that MITF knockdown affects melanoma cell proliferation(1–5). Hence, we investigated if MITF knockdown affects cell proliferation in U2OS cells. We used CellTrace^TM^ Violet dye to trace cell proliferation in MITF depleted cells. We stained cells with the CellTrace^TM^ dye prior to siRNA treatment that dilutes with each cell division. In this experiment the intensity of the cell dye was approx. 2.5-fold stronger in the MITF knockdown cells compared to a negative control, indicating reduced number of cell divisions after MITF knockdown (Figure 2E). Taken together, these results support that reduced MITF expression in U2OS cells causes chromosomal instability and negatively affects DNA replication and cell proliferation.

### MITF knockdown in U2OS cells accumulate replication stress and are more sensitive to chemotherapeutic drugs

Halted DNA replication can cause replication stress and urDNA when cells enter mitosis, which can potentially lead to the formation of bulky anaphase bridges (BABs)(15). As the cell divides, the enmeshed DNA can then give rise to DNA DSBs in the subsequent G1 phase, this damage forms structures known as 53BP1 nuclear bodies (NBs)(36, 37). To assess the impact of MITF loss in replication stress in U2OS cells, we used automated image analysis to quantify the number of 53BP1 foci in G1 phase (cyclin A negative). We found an increase in both BABs and 53BP1 NBs in MITF knockdown U2OS cells, indicative of increased replication stress after MITF loss (Figure 3A, B). To eliminate possible siRNA off-target effects, we repeated our experiment in the 624mel cell line expressing a dox-inducible MITF micro-RNA construct, (38). Inducible MITF knockdown revealed similar results as seen with RNAi mediated knockdown in U2OS, showing that MITF expression also suppresses genome instability in a melanoma cell line (Figure 3C and Supplementary Figure S3A).

**Figure 3.**
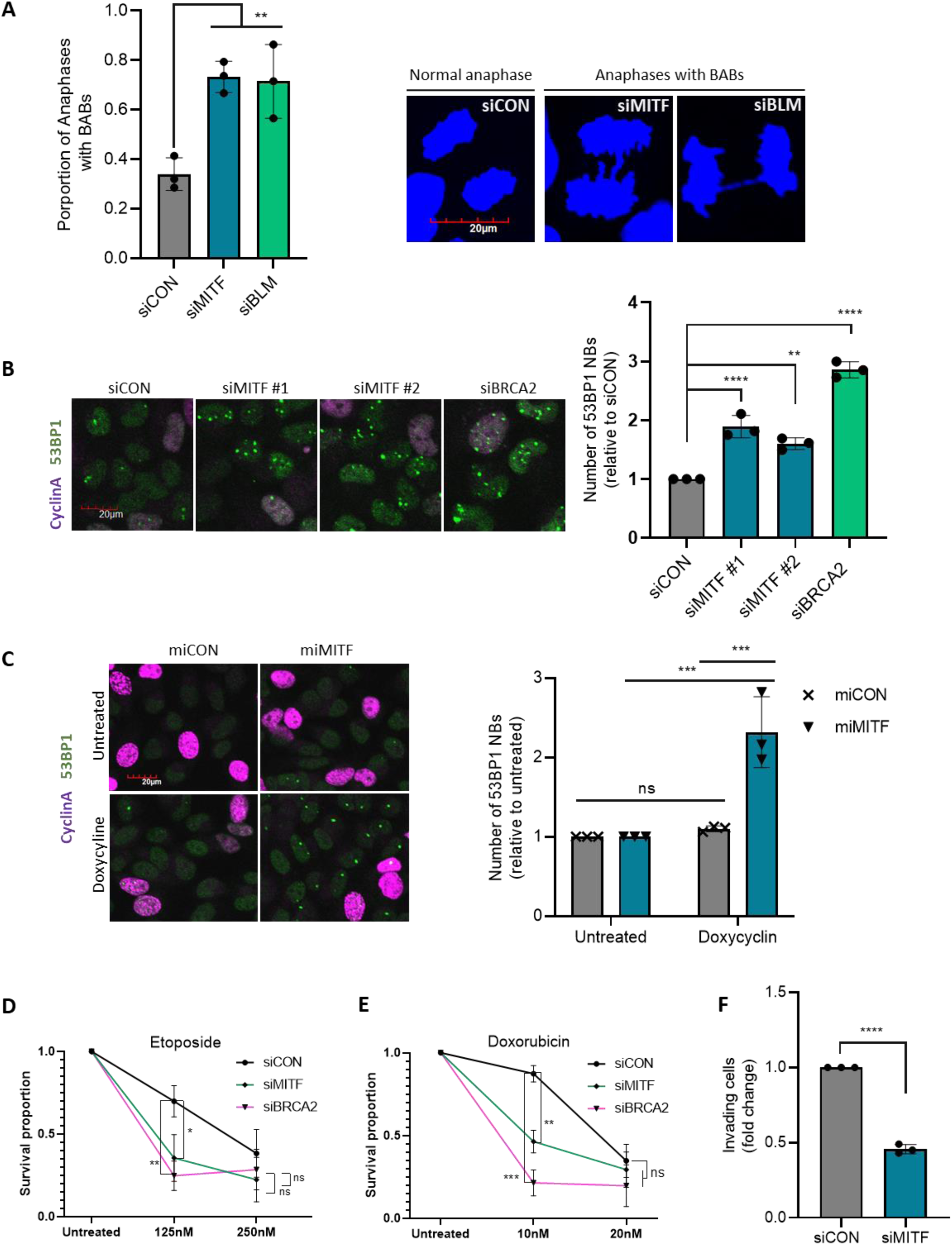
MITF knockdown in U2OS cells leads to replication stress and increased drug sensitivity. **a** Quantification (left) and representative images (right) of BABs after 72-hour siRNA treatment (siCON, siMITF and siBLM) followed by DAPI staining. Bar plot shows proportion of anaphase cells with BABs. Scale: 20μM (one-way-ANOVA, n=3). **b** Quantification and representative images of 53BP1 nuclear bodies after 48h siRNA treatment (siCON, siMITF, siBRCA1, si53BP1), followed by immunostaining with 53BP1 specific antibody. Scale: 20μM (one-way-ANOVA, n=3). **c** Quantification and representative images of 53BP1 nuclear bodies after 48h Doxycycline inducible MITF knockdown in 624mel cells, followed by immunostaining with antibodies targeting 53BP1 and CyclinA. Scale: 20μM (one-way-ANOVA, n=3). **d** Survival assay after seven-day siRNA treatment (siCON, siMITF and siBRCA2 as positive control) and Etoposide treatment. Cell survival was obtained with live-cell imaging by measuring cell confluency. The graph shows proportion of surviving cells compared to untreated samples (unpaired t-test, n=4). **e** Survival assay after seven-day siRNA treatment (siCON, siMITF and siBRCA2) and Doxorubicin treatment. Survival data presented as in d (unpaired t-test, n=3). **f** Matrigel invasion assay after 48h siRNA treatment (siCON and siMITF). Number of invading cells were obtained with EVOS FL fluorescent microscope following DAPI staining (unpaired t-test, n=3). Data presented as mean ± SD. \**P* <0.05, \*\**P* <0.01, \*\*\**P* <0.001, \*\*\*\**P* <0.0001.

Genome instability is in many cases associated with increased sensitivity to genotoxic agents. To test if increased chemotherapeutic drug sensitivity was observed after MITF knockdown, we used live-cell imaging to measure cell survival following drug treatment. We tested sensitivity to the topoisomerase II inhibitors doxorubicin and etoposide in U2OS cells, which are both drugs used in the treatment of various cancers, including osteosarcomas(39). We found that cell survival was lower after etoposide (Figure 3D) and doxorubicin treatment (Figure 3E) in MITF knockdown cells, indicating that MITF expression is important for cell survival in response to commonly used chemotherapeutic agents. This result uncovers new drug vulnerabilities in osteosarcoma.

Osteosarcoma has high metastatic rates. Approximately 85% of osteosarcoma patients suffer from metastatic disease, with the highest prevalence of pulmonary metastasis(40, 41). To test if MITF influenced the invasiveness of U2OS cells, we performed Matrigel transwell invasion assay. Knockdown of MITF resulted in a clear reduction in number of invading cells (Figure 3F and Supplementary Figure S3B).

Collectively this data shows that upon MITF knockdown, U2OS cells have decreased invasiveness and are more sensitive to cancer drug exposure.

### MITF knockdown activates the P53 stress signaling pathway

P53 plays a key role in cell cycle regulation and apoptosis in response to DNA damage(42). Given the observed genome instability phenotypes, we decided to look at P53 activation before and after MITF depletion. Knockdown of MITF, using siRNAs, resulted in a two-fold increase in P53 protein levels and a significant increase in mRNA expression (Figure 4A, B). Consistent results were obtained in the melanoma cell line 624mel, following RNAi and doxycycline inducible knockdown of MITF (Supplementary Figure S4A, B, C). Furthermore, MITF knockdown in U2OS cells affected cell cycle distribution whereas increased proportion of cells were observed in G1-phase, indicating cell cycle arrest (Figure 4C). Importantly, in the isogenic U2OS P53 knock-out cell line(43) no G1 arrest is observed, supporting the that the cell cycle arrest is mediated by P53 (Supplementary Figure S4D).

**Figure 4.**
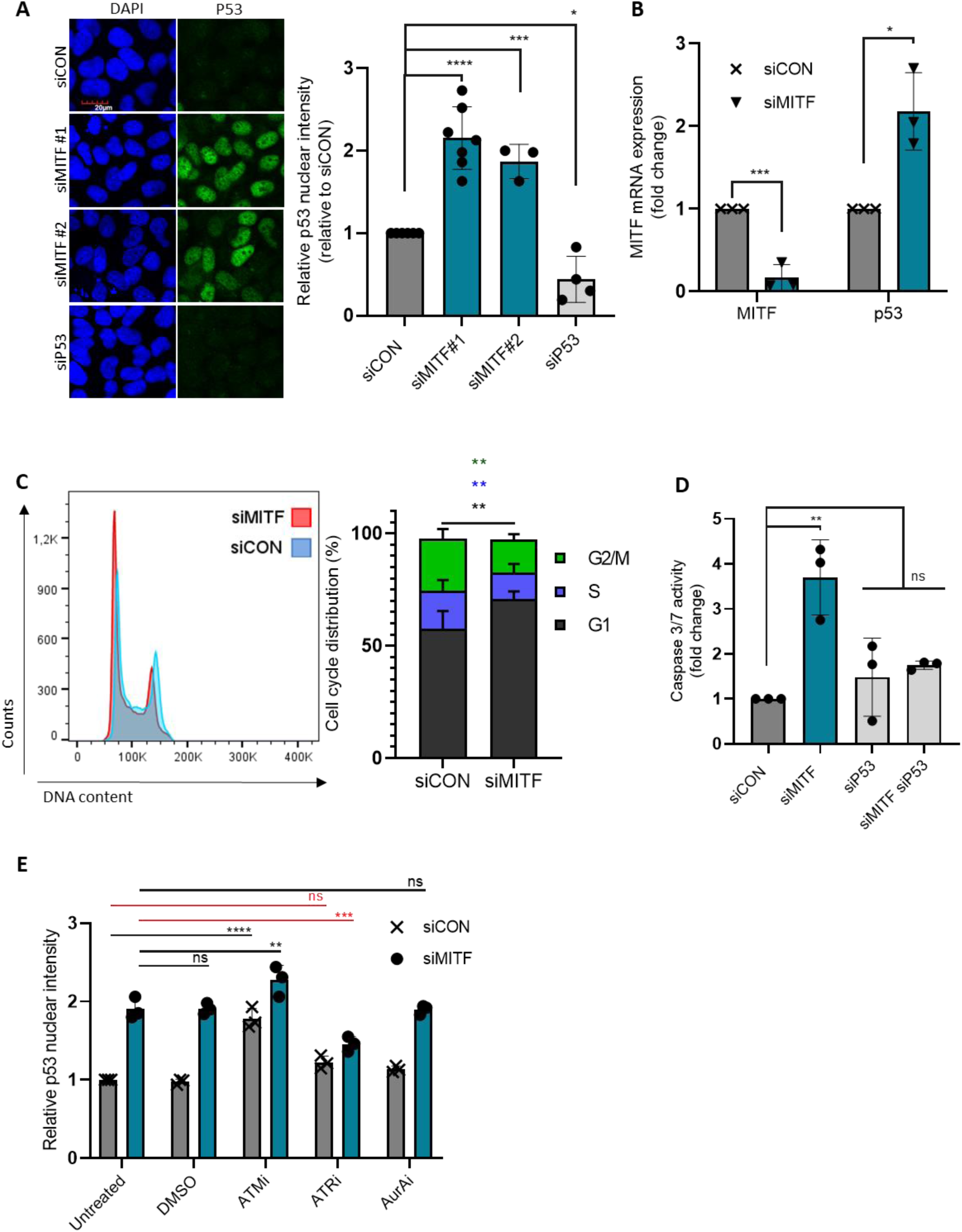
P53 is activated upon MITF knockdown in U2OS cells. **a** Confocal microscopy images and quantification of P53 protein levels after 48h treatment with siCON, siMITF and siP53, followed by immunostaining with antibody targeting P53 and DAPI to visualize nuclear cells. Scale: 20μM (one-way-ANOVA, n=3-7). **b** MITF and p53 mRNA expression in U2OS cells. Cells were treated with indicated siRNAs for 48h, followed by real time qPCR analysis. (unpaired t-test, n=3). **c** Cell cycle profile analyses of siCON and siMITF treated U2OS cells. Cells were fixed after 48h siRNA treatment, followed by staining of DNA content with 7- aminoactinomycin D (7AAD) and flow cytometry analysis (unpaired t-test, n=4). **d** Caspase 3/7 activity relative to cell confluency after four days of siRNA treatment (siCON, siMITF, sip53, siMITF + sip53). Cells were incubated with caspase-3/7 red dye after siRNA treatment and imaged in IncuCyte^®^ live-cell imager (one-way-ANOVA, n=3). **e** quantification of P53 protein levels after 48h siCON and siMITF treatment and 24h treatment with ATMi, ATRi, AuroraAi or DMSO, followed by immunostaining with antibody targeting P53 and DAPI to visualize nuclear cells (one-way ANOVA, n=3). Data presented as mean ± SD. \**P* <0.05, \*\**P* <0.01, \*\*\**P* <0.001, \*\*\*\**P* <0.0001.

To verify if MITF knockdown would affect the survival of the U2OS cell line, we next sought to evaluate if the cells would enter an apoptotic state upon treatment with MITF siRNA. Caspase cleavage assay revealed an increase in apoptosis after MITF knockdown, with an approx. 4-fold increase in caspase cleavage activity after 96 h siRNA treatment (Figure 4D). Collectively this data suggests that MITF depleted cells activate the P53 mediated stress response, resulting in cell cycle arrest and apoptosis. These results are also in line with the cell proliferation analysis, whereas reduced cell proliferation of MITF depleted cells is rescued with a double knockdown of MITF and P53 (Figure 2E).

To further characterize the upstream mechanism of the P53 stress response observed in MITF knockdown U2OS cells, we inhibited key members of the phosphoinositide-3-kinase-related protein kinase (PIKK) family, which play a central role in the DNA damage response. Inhibiting ATM and AuroraA had no effect on P53 accumulation following MITF knockdown. However, inhibition of ATR, a key kinase activated during replication stress, significantly reduced P53 accumulation, suggesting that problems during replication are involved in P53 activation in MITF depleted cells (Figure 4E).

Furthermore, given the observed relationship between MITF knockdown and p53 expression, we used TCGA to quantify the relationship in expression between these two genes. Out of 32 different cancer types, 18 showed negative correlation between MITF and p53 (sarcoma shown as an example, Supplementary figure S4E).

### MITF knockdown promotes P53 activation and cell cycle arrest through LATS2

Upon cellular stress, P53 is activated by tightly regulated protein modifications, which rapidly results in P53 protein accumulation and subsequent cell cycle arrest. P53 is phosphorylated at various sites, causing conformational changes, which blocks ubiquitination-mediated degradation by MDM2(44–46). It has also been shown that in response to mitotic stress and chromosomal instability, LATS2 is translocated to the nucleus where it binds MDM2 and inhibits its ligase activity towards P53, resulting in rapid P53 accumulation(21, 23, 28, 29). In this context, P53 enhances LATS2 mRNA expression, creating a positive feedback loop that results in strong P53 accumulation(21, 23). As we shown above, MITF knockdown yields a dramatic increase in P53 expression and increased number of chromosome aberrations (Figures 4A and 2A). Therefore, we investigated if P53 activation in MITF depleted cells is dependent on LATS2 activity. To this end, we did double knockdowns of MITF and LATS2 in U2OS cells to examine the impact on P53 expression and cell cycle arrest. We found that co-depletion of MITF and LATS2 resulted in a complete rescue of both P53 protein accumulation and cell cycle arrest (Figure 5A, B and Supplementary Figure S5A, D), suggesting strong dependency on LATS2 of the Hippo pathway. Additionally, co-depletion of MITF and LATS2 in 624mel cells partially rescued the P53 protein accumulation (Supplementary Figure S5B). Attempting to rescue G1 cell cycle arrest in the 624mel cell line was not possible since the cells are not arrested in the G1 phase upon MITF depletion (Supplementary Figure S5C). This is most likely due to a mutation in the DNA binding domain of P53 in 624mel cells (https://www.cellosaurus.org/CVCL_8054). To confirm LATS2 activation, we analyzed nuclear translocation of Myc-LATS2 in control and MITF knockdown cells. This revealed a 3- fold increase of LATS2 in the nucleus in MITF depleted cells, indicating LATS2 activation (Figure 5C). qPCR analysis revealed up-regulation of LATS2 mRNA after MITF knockdown (Figure 5D), suggesting that LATS2 is also under MITF control at the transcription level.

**Figure 5.**
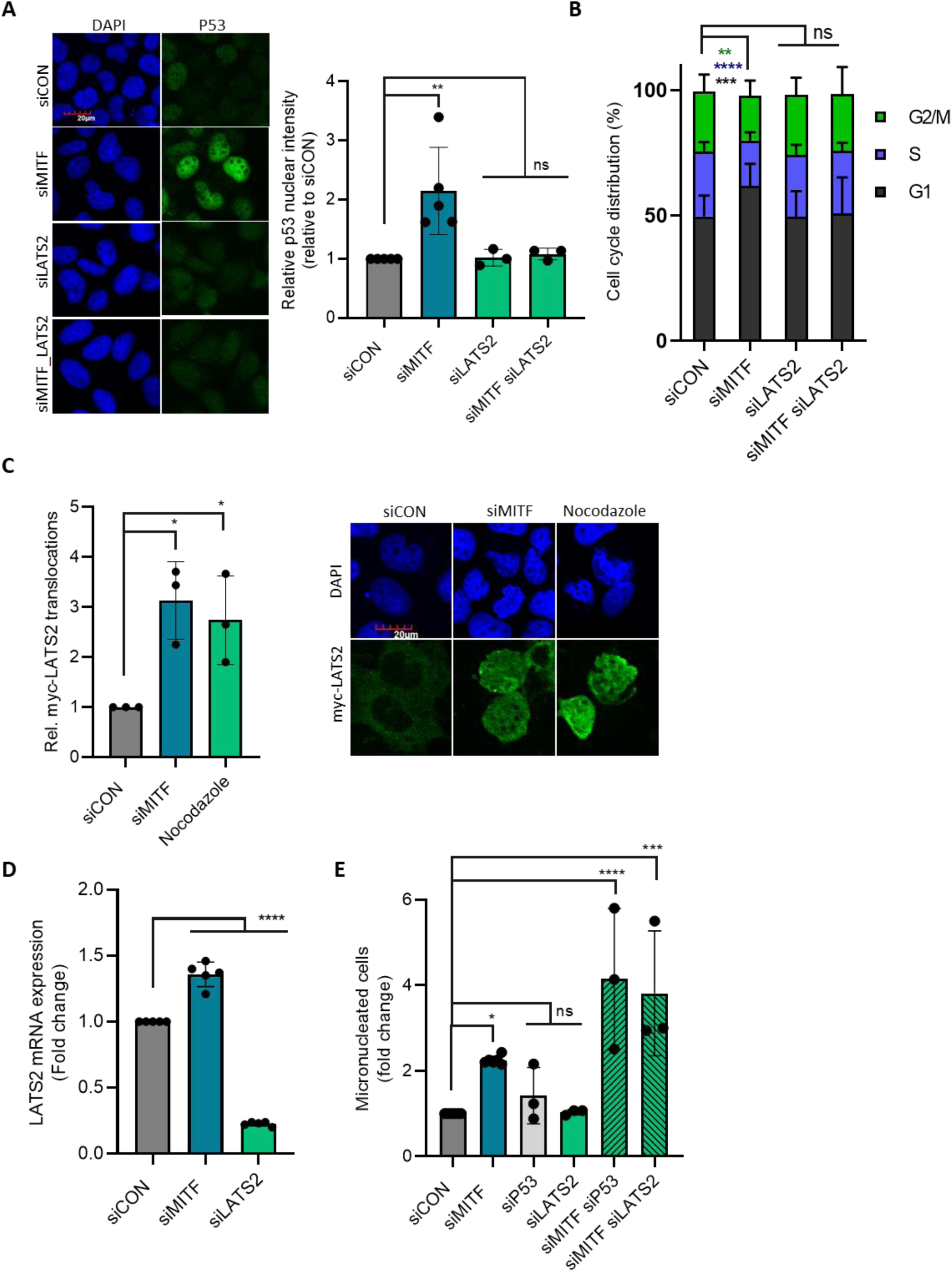
P53 activation in MITF knockdown cells is dependent on the Hippo pathway kinase LATS2. **a** Confocal microscopy images (left) and quantification (right) of P53 protein levels in U2OS cells 48h after treatment with siCON, siMITF, siLATS2 and siMITF + siLATS2, cells were fixed and immunostained with a P53 specific antibody and DAPI to visualize nuclear cells. Scale: 20μM (one-way-ANOVA, n=3-5). **b** Cell cycle profile analyses of siCON, siMITF, siLATS2 and siMITF + siLATS2 treated U2OS cells. Cells were fixed after 48h siRNA treatment, followed by staining of DNA content with 7-aminoactinomycin D (7AAD) and flow cytometry analysis (one-way-ANOVA, n=4). **c** Confocal images (right) and quantification (left) of Myc-LATS2 translocation into the nucleus in siCON and siMITF treated U2OS cells. 24h Nocodazole treatment was used as a positive control. Cells were treated with siRNA for 48h, Myc-LATS2 plasmid transfection was performed 24h prior to fixation. This was followed by immunostaining with Myc-tag specific antibody and DAPI to visualize nuclear cells. Scale: 20μM (one-way-ANOVA, n=3). **d** MITF and LATS2 mRNA expression in U2OS cells after 48h siCON and siMITF treatment. mRNA levels were analyzed using qPCR (one-way-ANOVA, n=4). **e** Quantification of the proportion of cells containing micronuclei following 72h siRNA treatment (siCON, siMITF, siP53, siLATS2, siMITF + siP53 and siMITF + siLATS2) and DAPI staining. The bar plot shows fold change in cells with micronuclei compared to siCON (one-way-ANOVA, n=3). Data presented as mean ± SD. \**P* <0.05, \*\**P* <0.01, \*\*\**P* <0.001, \*\*\*\**P* <0.0001.

As a key activator of DNA damage checkpoint signaling, P53 plays an important role in preventing replication in damaged cells and subsequent loss of genetic integrity. Accordingly, we observed increased genome instability after MITF knockdown, when P53 activation was inhibited by co-depleting the cells with siLATS2 or siP53 (Figure 5E and Supplementary Figure 5SD, E). Furthermore, we found that LATS2 or P53 knockdown alone had no effect on genome instability, it only affects the cells in combination with MITF depletion. This supports that LATS2 dependent activation of P53 plays a key role in counteracting genome instability caused by reduced MITF expression.

### MITF expression levels are clinically relevant in melanoma and non-melanoma cancer

MITF has a well-established role as an oncogene in melanoma. Indeed, increased MITF expression has been associated with poor prognosis in melanoma patients(6). Therefore, we sought to investigate the effect of MITF expression in patient survival in non-melanoma cancers. Using data from the Cancer Genome Atlas (TCGA), we generated Kaplan-Meier plots in patient cohorts separated by MITF gene expression levels. MITF expression associates with survival in sarcoma, adrenocortical carcinoma and kidney clear cell carcinoma as well as melanoma (Supplementary Figure 6). This shows that changes in MITF expression are clinically relevant in melanoma and non-melanoma cancers. Conclusively, low MITF expression associates with poor survival in the non-melanoma cancer types, which may be indicative of a tumor suppressive role of MITF.

## Discussion

Despite a well-illustrated role of MITF in melanocytes and melanoma, it has been shown that MITF has a role in osteoclast and mast cell development, olfactory bulb function, autophagy, and retinal pigment epithelium (RPE) development(7, 47–53). In recent years, MITF has also been suggested to have melanocyte-specific roles in mitotic regulation, DNA replication and DNA damage repair, all of which are important to maintain genome integrity(16–18). Here, we show that MITF has an important function in maintaining genome stability in cell lines derived from osteosarcoma (U2OS), chondrosarcoma (SW-1353), cervical cancer (HeLa), and the non-cancer breast epithelia cell line D492. This suggests that MITF preserves genome integrity in a variety of tissue types and is not limited to cells of melanocytic origin. We show that non-melanocyte cell lines that express low levels of MITF compared to melanoma cell lines depend on MITF to maintain their genome integrity (Figure 1D, E and Supplementary Figure S1A-D).

Genome instability is considered as one of the hallmarks of cancer and a major driving force for tumorigenesis. In light of that, our results indicate that in addition to its established role as a melanoma specific oncogene, MITF might also play a broader role as a genome maintenance factor, which can potentially contribute to the recently described tumor suppressive function of MITF(54).

Although our results clearly show that MITF is important for maintaining genome stability in various tissue types, there might be examples where other mechanisms are at play. One out of the six cell lines tested, A549, did not show any signs of genome instability following MITF knockdown (Figure 1D, E). Mechanistically we do not understand the lack of phenotype in A549. Potentially, A549 harbors a mutation or an epigenetic modification, which neutralizes the damaging effect of MITF depletion. Nevertheless, these results are interesting, since they show that there is a level of tissue specificity involved in MITF’s function in genome maintenance, which merits further research.

We also show that MITF knockdown negatively affects cell proliferation of U2OS cells and that long-term MITF depletion ultimately results in apoptosis (Figures 2D and 4D). Our findings show an impairment in cell cycle progression, with less cells in S-phase and slower incorporation of nucleotides upon MITF knockdown in U2OS cells (Figure 2C, D 4C and Supplementary table S1 and S2), indicating decreased DNA replication. Markedly it has been reported that DNA replication is under MITF regulation in melanoma cells(18). Moreover, our data suggests that the stress response observed in MITF knockdown cells is mediated through ATR and LATS2 of the Hippo pathway (Figure 4E and Figure 5A-D), revealing a novel connection between MITF and key signaling factors in the response to DNA damage. From a mechanistic perspective, the connection with ATR gives important insight into the source of genome instability in MITF deficient cells. The ATR kinase is activated at stressed replication forks and plays a key role in maintaining the integrity DNA replication under stress conditions(55, 56) . The need for ATR in LAT2/P53 stress signaling further supports that replication stress is a main source of genome instability in MITF deficient cells.

Previously, it has been shown that MITF directly regulates genes required for DNA replication in melanoma cells(17, 18, 57). This is in accordance with our data, where DNA replication pathways scored as significantly enriched terms in the set of downregulated genes in U2OS MITF knockdown cells. Interestingly, major DNA repair pathways were also significantly enriched, pointing to a potential mechanism for impaired cell replication (Figure 2C and Supplementary table S2).

As expected, our data revealed expression of both the A-and the M-isoform in the melanocytic SkMe28 cell line. Interestingly, the U2OS cell line expresses the A- and the H-isoforms, whereas the A-isoform is predominantly expressed (Figure 1G). This raises the question whether different MITF isoforms carry out different functions. Majority of studies on MITF in melanoma or melanocytes do not differentiate between isoforms. However, in a recent study, an isoform-specific function of the A-isoform as a transcriptional regulator of genes involved in autophagy and oxidative metabolism was observed (58). Furthermore, a study using isoform-specific knockout mice reveals that the M-isoform specifically contributes to pigmentation, while knockout of the A-isoform had only a minor effect on that phenotype(59, 60). The phenotype reported here is most likely designated to the A-isoform as it is the MITF isoform predominantly expressed in the U2OS cell line, however, this does not exclude the possibility of other MITF isoforms participating in genome stability.

Melanoma is an aggressive cancer type with high metastatic potential and poor 5-year survival rates(6). MITF is proposed to act as a rheostat, by controlling the switch between different signaling pathways that determine the phenotype of melanoma cells(61) making melanoma frequently resistant to therapeutic treatments(4, 5, 62). According to the rheostat model, high MITF levels in melanoma cells are linked to proliferating phenotype, while cells expressing lower levels of the protein are less proliferative and more invasive and MITF depletion ultimately results in cell death. However, more recent studies have not been able to replicate the more invasive phenotype in *in vitro* studies(38, 63). In these studies, MITF knockdown in melanoma cells has little to no effect on invasion ability, but knockout results in less invasion ability. Our data suggests a significant reduction in invasive potential (Figure 3F), but like melanoma, osteosarcoma has high metastatic rates and 5-year survival of metastatic patients as low as 20%(40, 41). It should be noted that the tumor microenvironment plays an important role in tumor cell invasion(62) which our experimental settings are not able to simulate. Therefore, the reduced ability to invade through the Matrigel might be a consequence of overall reduced fitness of the MITF knockdown cells.

Results from our cell line analysis (Figure 1) indicate MITF has an important function in melanoma and non-melanoma cancers. In line with that, we show that MITF expression has a significant effect on survival in various non-melanoma cancers (Supplementary Figure S6). This supports our hypothesis and underscores the importance of characterizing the role of MITF across cell lineage. Furthermore, results from our drug sensitivity experiments revealed that MITF knockdown increases sensitivity of U2OS cells to the chemotherapeutic drugs, Etoposide and Doxorubicin (Figure 3D, E)(39), which indicates that MITF levels might determine treatment outcomes not only in melanoma patients but in other cancer patients as well.

As transcription factor, MITF regulates the expression of a large number of genes, with approximately 500 genes as validated targets(64). Functionally, MITF target genes are quite diverse including genes involved in pigmentation, DNA damage response, mitotic regulation and DNA replication(18, 64). Our data suggests that the role of MITF as a regulator of genome maintenance extends beyond the melanocyte cell lineage by regulating expression of genes with roles in DNA replication and DNA repair (Supplementary tables S1 and S2). These transcriptional targets of MITF have been reported in melanoma cells on many occasions in the literature(16–18), but here we propose that this function is not melanocyte specific, suggesting MITF expression is biologically relevant in other tissue types. Dysregulation of LATS2 expression and the Hippo pathway, has been linked to tumorigenesis and cancer progression in many cancer types, (24–27, 65–68). Here we show a novel connection between MITF and LATS2, a key kinase in the Hippo pathway, which in light of its important role in tumor progression, further supports the potential tumor suppressive role of MITF.

## Data availability statement

Flow cytometry data has been deposited in the Flow depository: http://flowrepository.org/id/RvFrVYTvPpVgLRpylPZjOj0zanKPIEzRCG7LUrZMTm5MQWkC 0Xzt2aXukiZpFutV (ID: FR-FCM-Z7B8) and http://flowrepository.org/id/RvFrqyXBg2JWNta8wrUFC7dAdozDqQaOo50of0pveXDjvZn40 UeKytWhR8HRwUuF (ID: FR-FCM-Z7CG).

RNAseq data has been deposited in the GEO depository: https://www.ncbi.nlm.nih.gov/geo/info/update.html (private access token: kjwxgoywjpotxwb)

## Acknowledgements

We would like to thank the Functional Genetics Laboratory at DeCode Genetics Reykjavik Iceland for generously providing use of their Attune^TM^ NxT machine. We also thank the Landspitali University Hospital for providing use of their Beckman Coulter Navios Ex flow cytometer. The P53-KO U2OS cell line was graciously provided by Libor Macurek. We would also like to thank The Icelandic Cancer Society research fund and The Icelandic research fund (RANNIS) for their financial support.

## Contributions

TG and StS were responsible for study design. DHG contributed to the study design, performed confocal imaging assays, Flow cytometry assays, Matrigel trans well assays and live-cell imaging assays along with manuscript writing. TG, StS, LV and ALGM contributed to manuscript writing. RNA sequencing and analysis was performed by SnS, KK, ALGM and DM. Isoform analysis was performed by KK.. ESv and SR performed the metaphase spreads and aided in the analysis of chromosome images. Doxycycline inducible cell lines were designed and made by RD. TG performed WB analyses and TBV and MRB performed qPCR analyses. Data interpretation was done by DHG, TG, StS and ALGM. ESt contributed intellectually to the study and writing of the manuscript.

