## Supplementary Figures for "MITF maintains genome stability in non-melanocytic cell lineages and suppresses Hippo pathway signaling"

### Figure s1

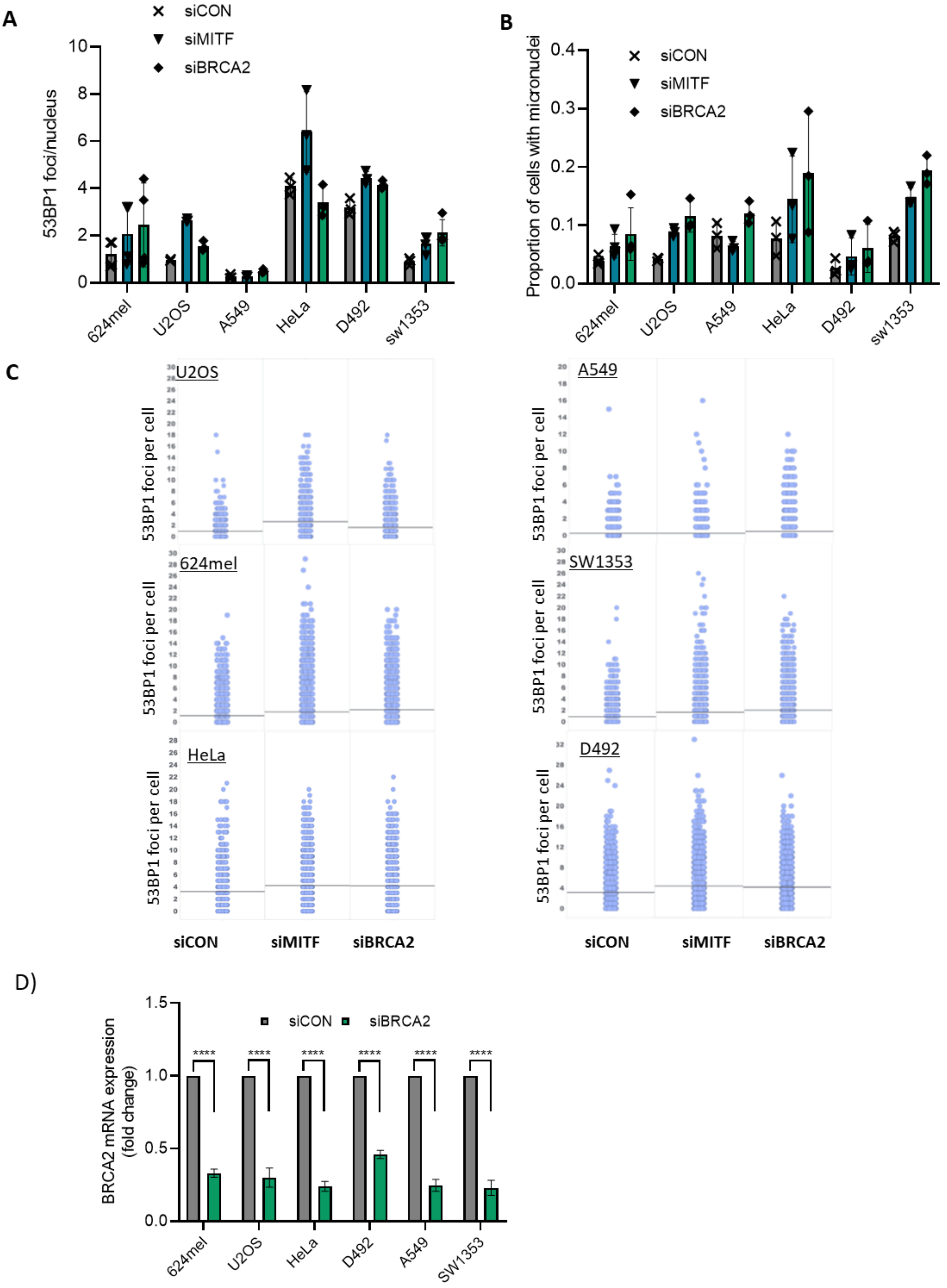
A
siCON
siMITF
siBRCA2
53BP1 foci/nucleus
624mel
U2OS
A549
HeLa
D492
sw1353
B
Proportion of cells with micronuclei
C
U2OS
A549
624mel
SW1353
HeLa
D492
53BP1 foci per cell
siCON
siMITF
siBRCA2
D)
BRCA2 mRNA expression (fold change)
SW1353
****

**Supplementary Figure S1 A** Graphs representing mean of actual numbers of nuclear 53BP1 foci in 624mel, U2OS, A549, HeLa, D492 and SW1353 following treatment with the indicated siRNAs for 72h and immunostaining with 53BP1 specific antibody. **B** Graphs representing proportion of cells with micronuclei in 624mel, U2OS, A549, HeLa, D492 and SW1353 following treatment with the indicated siRNAs for 72h and staining with DAPI nuclear stain. **C** 53BP1 foci distribution plot from the indicated cell lines following 48h treatment with control, MITF and BRCA2 targeting siRNAs. Each dot represent an individual cell, black horizontal line show the average number of foci from three independent experiments. **D** BRCA2 mRNA expression in indicated cell lines. Cells were treated with siRNA targeting BRCA2 or siRNA control for 48h, followed by real time qPCR analysis. (unpaired t-test, n=3).

Figure s2

A

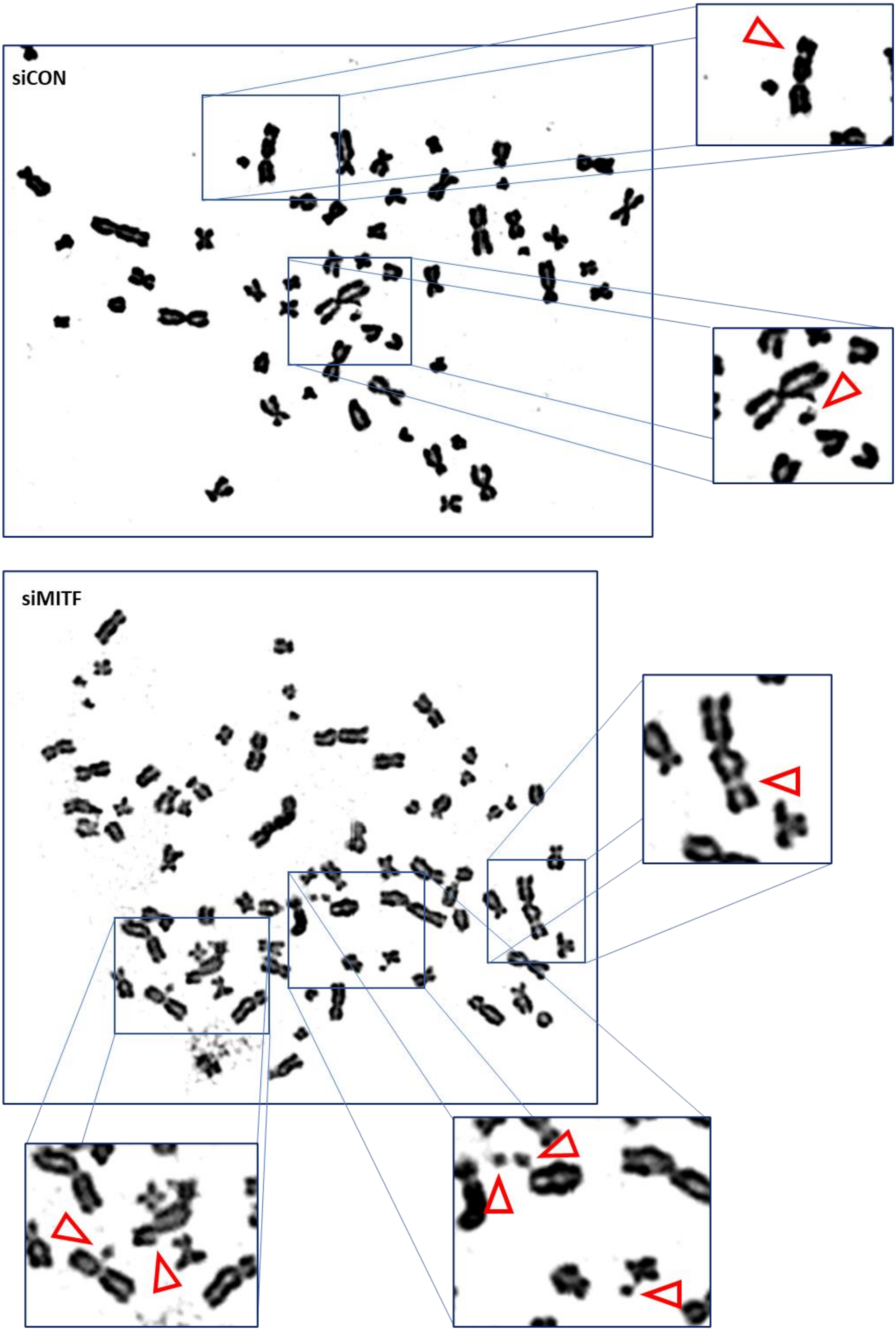

**B**

**Differentially expressed genes between MITF KD and WT conditions in SkMel28**

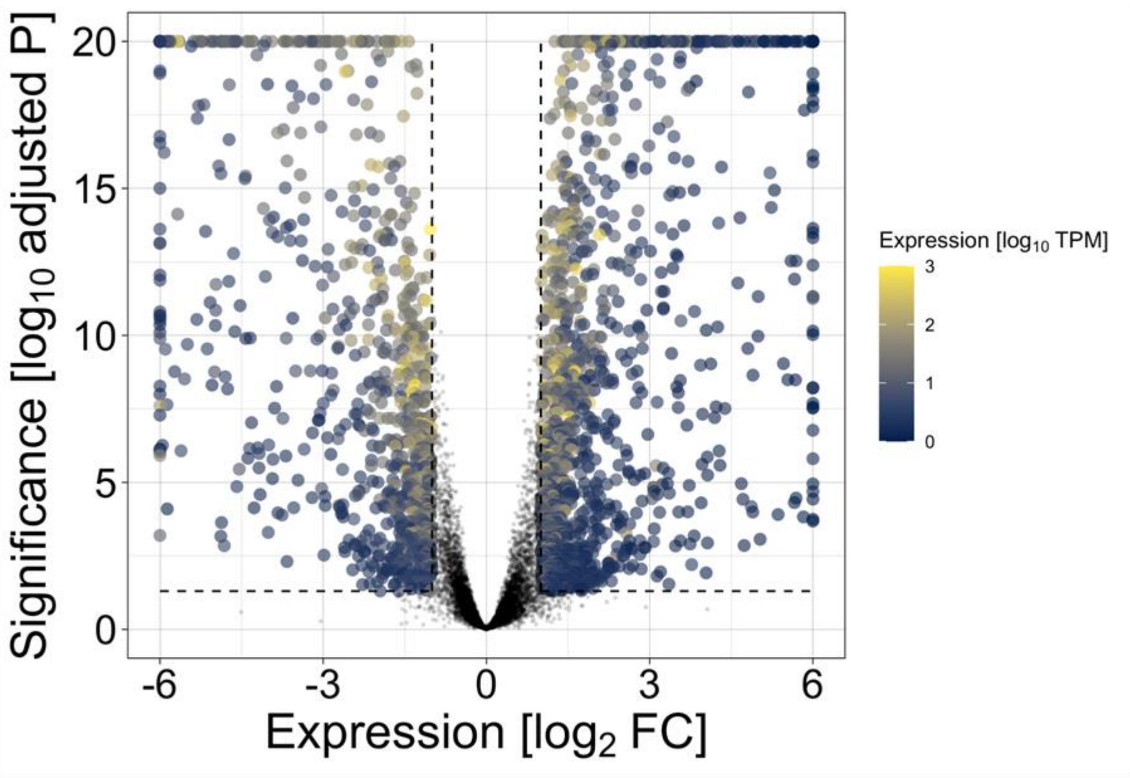


**Differentially expressed genes between MITF KD and WT conditions in U2OS**

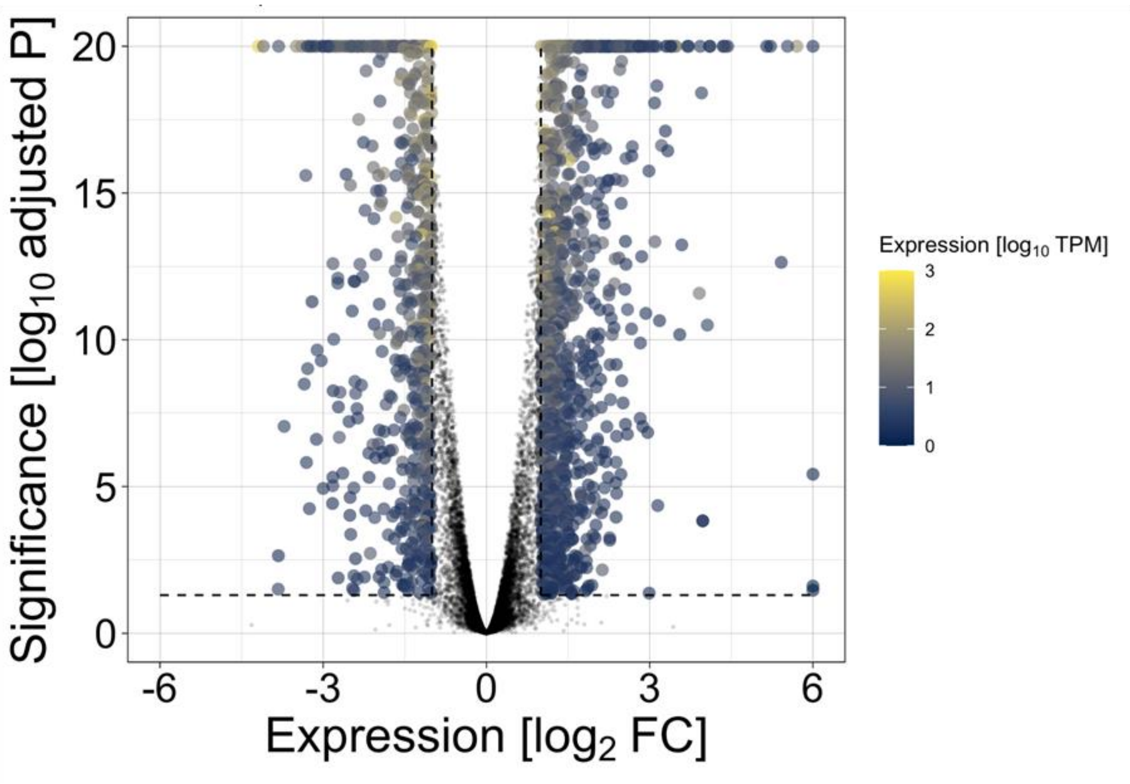

**Supplementary Figure S2 A** Representative images of metaphase spreads for siCON (upper panel) and siMITF (lower panel) treated cells. Cells were treated with siRNA for seven days. To enrich cells in metaphase, samples were treated with the mitotic inhibitor Colcemid four hours prior to fixation. **B** Volcano plots showing number of genes affected by 48h siRNA mediated MITF knockdown in SkMel28 cells (upper panel) and U2OS cells (lower panel). Each dot in the volcano plot represents a gene affected by MITF knockdown in U2OS cells. Genes that are non-significantly affected are represented in black dots.

### Figure s3

A

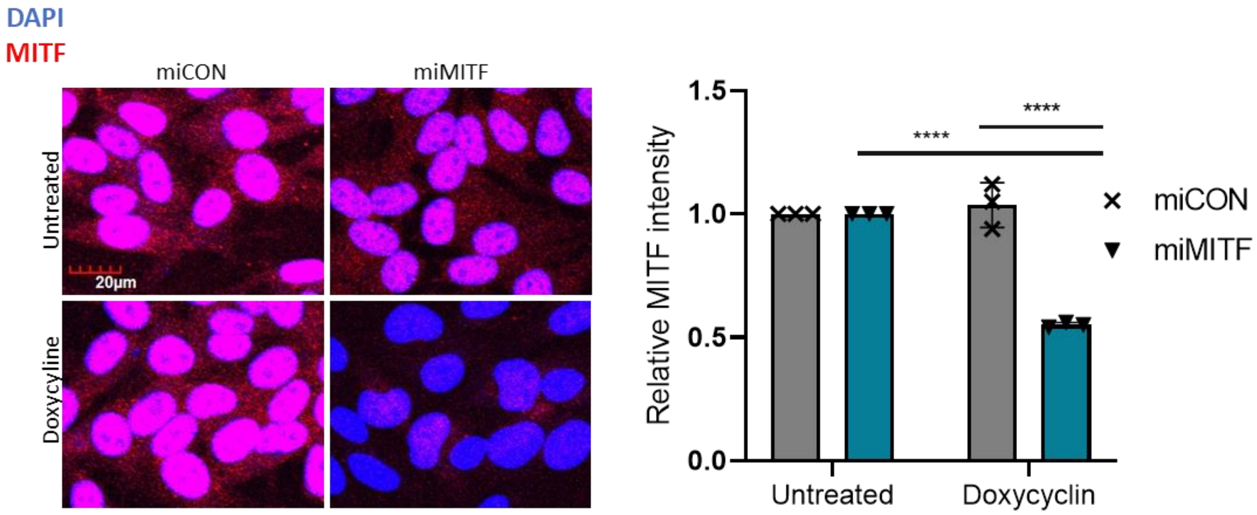


B

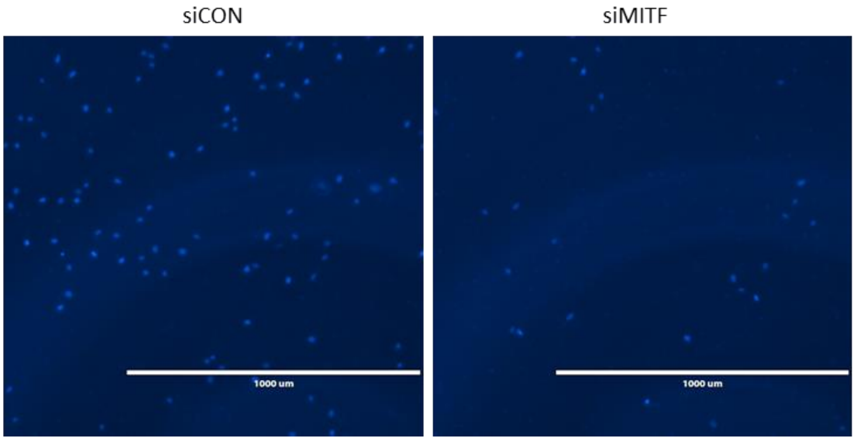


**Supplementary Figure S3 A** Quantification (right) and representative images (left) of MITF expression before and after 48 Doxycycline inducible MITF knockdown in 624mel cells. This was followed by immunostaining with MITF antibody (one-way ANOVA, n=3). **B** Representative fluorescent microscope images showing invading cells after 48h siRNA treatment (siCON and siMITF), followed by DAPI staining. Data presented as mean ± SD. **P* <0,05, ***P* <0,01, ****P* <0,001, *****P* <0,0001.

### Figure s4

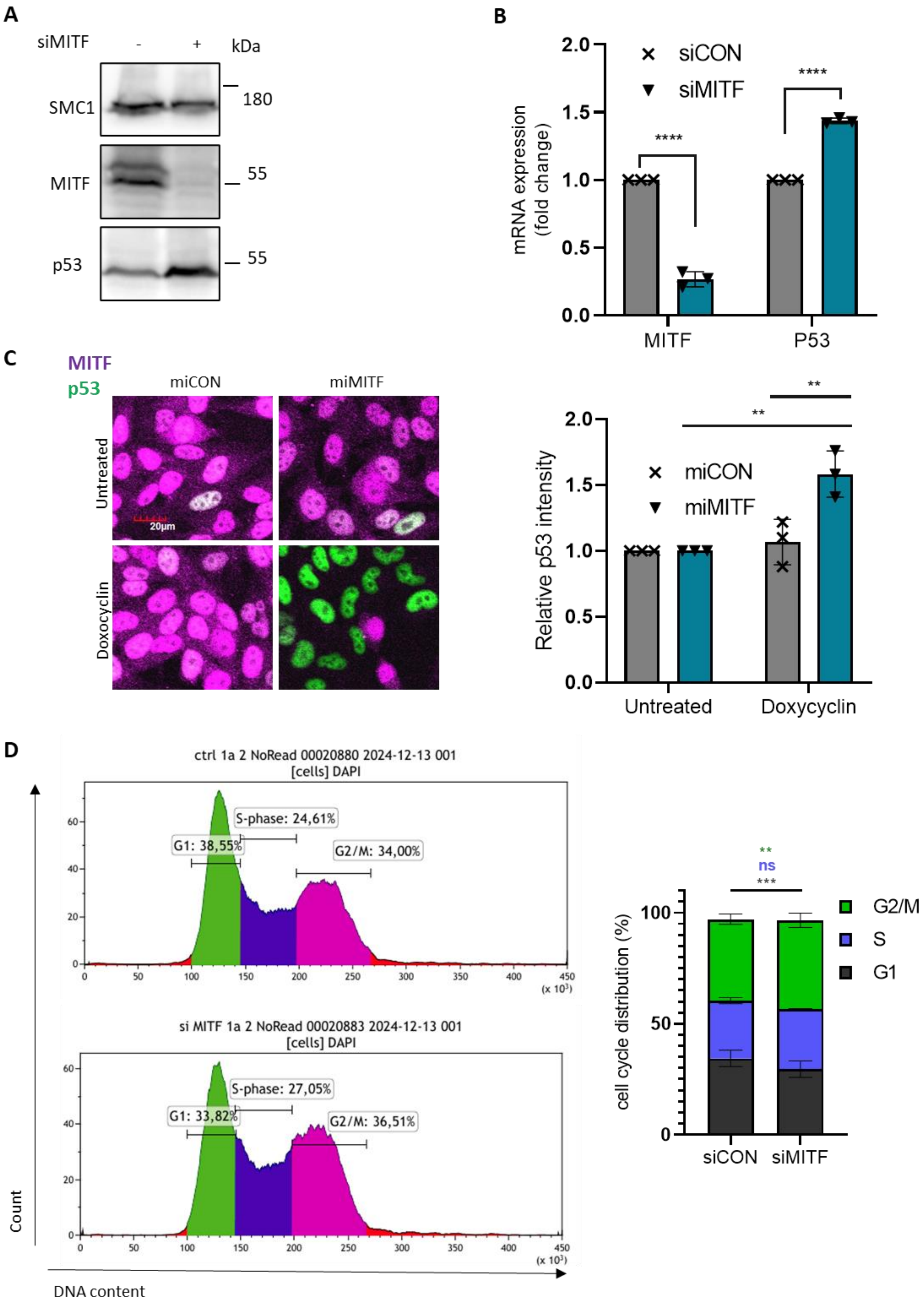
A
siMITF
-
+
kDa
SMC1
180
MITF
55
p53
55
B
mRNA expression (fold change)
siCON
siMITF
****
****
MITF
P53
C
MITF
p53
miCON
miMITF
Untreated
Doxocyclin
20µm
Relative p53 intensity
miCON
miMITF
**
**
Untreated
Doxycyclin
D
ctrl 1a 2 NoRead 00020880 2024-12-13 001
[cells] DAPI
S-phase: 24,61%
G1: 38,55%
G2/M: 34,00%
si MITF 1a 2 NoRead 00020883 2024-12-13 001
[cells] DAPI
S-phase: 27,05%
G1: 33,82%
G2/M: 36,51%
Count
DNA content
cell cycle distribution (%)
**
ns
***
G2/M
S
G1
siCON
siMITF

**E**

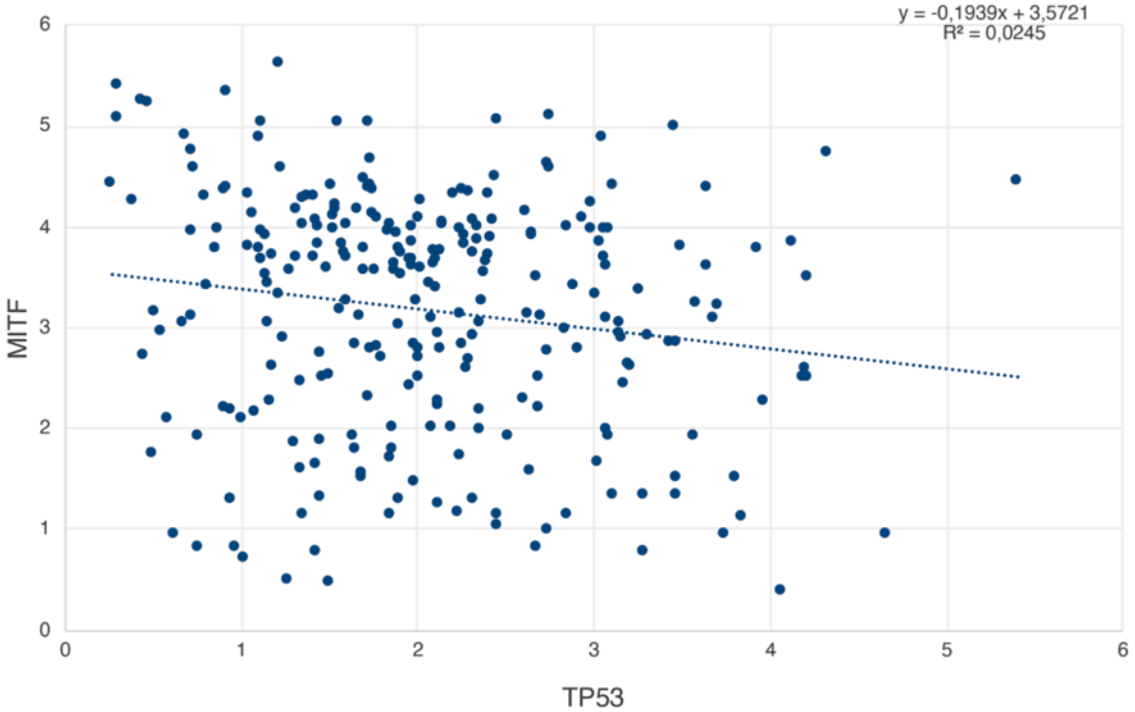


**Supplementary Figure S4 A** MITF and P53 protein expression in 624mel cells after 48h siRNA treatment (siCON and siMITF), analyzed by western blot of whole cell extracts using indicated antibodies. SMC1 was used as loading control. **B** MITF and P53 mRNA expression in 624mel cells. Cells were treated with indicated siRNAs for 48h, followed by real time qPCR analysis (unpaired t-test, n=3). **C** Quantification (right) and representative images (left) of P53 protein expression before and after 48h Doxycycline inducible MITF knockdown in 624mel cells. This was followed by immunostaining with MITF and P53 antibodies (one-way ANOVA, n=3). **D** Cell cycle profiles and graphs of siCON and siMITF treated U2OS-stable P53 knockout cells. Cells were fixed after 48h siRNA treatment, followed by staining of DNA content with DAPI nuclear stain and flow cytometry analysis (unpaired t-test, n=3). **E** Graph showing negative correlation between the expression of MITF and P53 in sarcoma patient samples. Each dot on the graph represents MITF expression (y-axis) and P53 expression (x-axis) of single tumor. Values were extracted from the TCGA database Data presented as mean ± SD. **P* <0,05, ***P* <0,01, ****P* <0,001, *****P* <0,0001.

### Figure s5

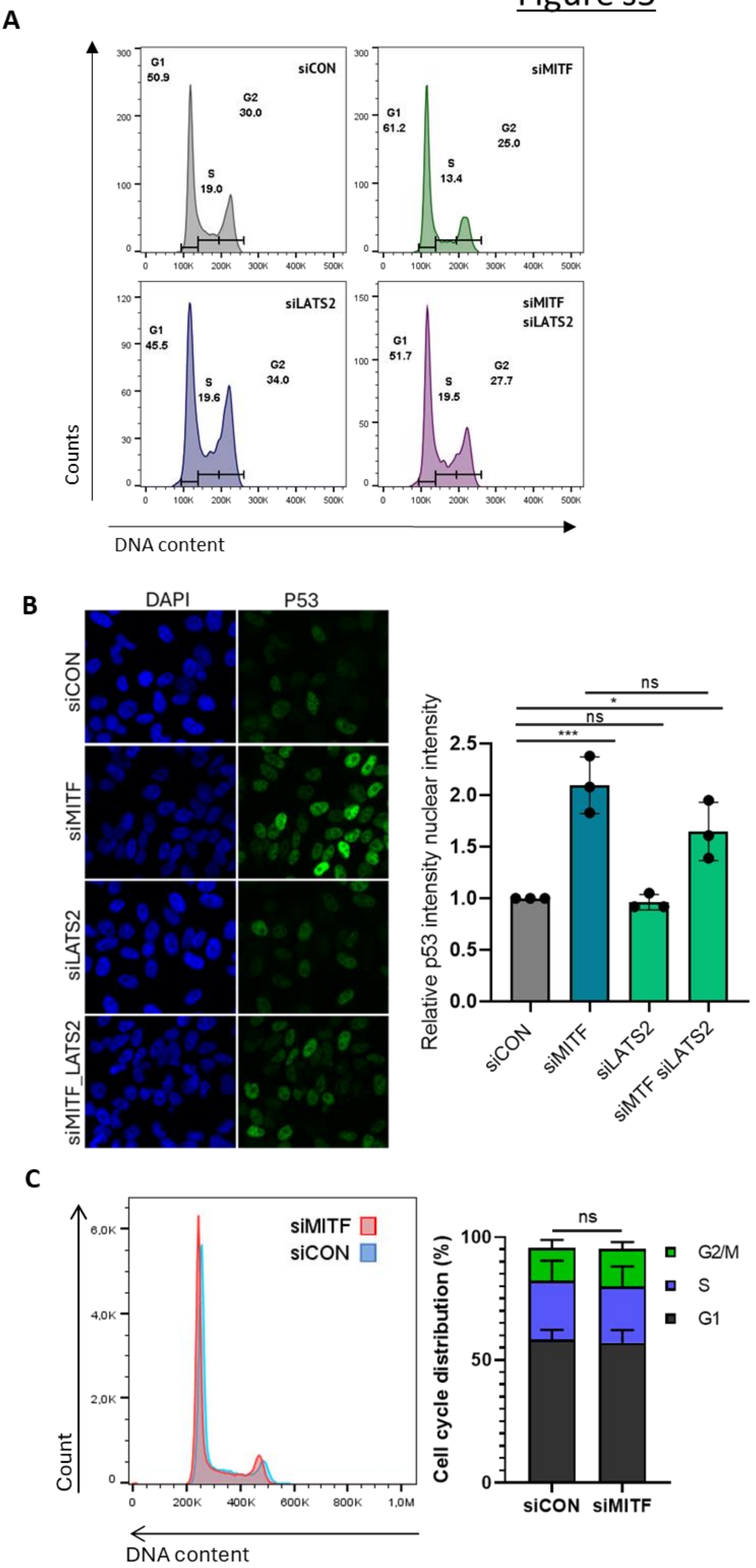
A
siCON
G1
50.9
S
19.0
G2
30.0
siMITF
G1
61.2
S
13.4
G2
25.0
siLATS2
G1
45.5
S
19.6
G2
34.0
siMITF
siLATS2
G1
51.7
S
19.5
G2
27.7
Counts
DNA content
B
DAPI
P53
siCON
siMITF
siLATS2
siMITF_LATS2
Relative p53 intensity nuclear intensity
ns
*
ns
***
siCON
siMITF
siLATS2
siMTF siLATS2
C
siMITF
siCON
Count
DNA content
ns
Cell cycle distribution (%)
G2/M
S
G1
siCON
siMITF

**D**

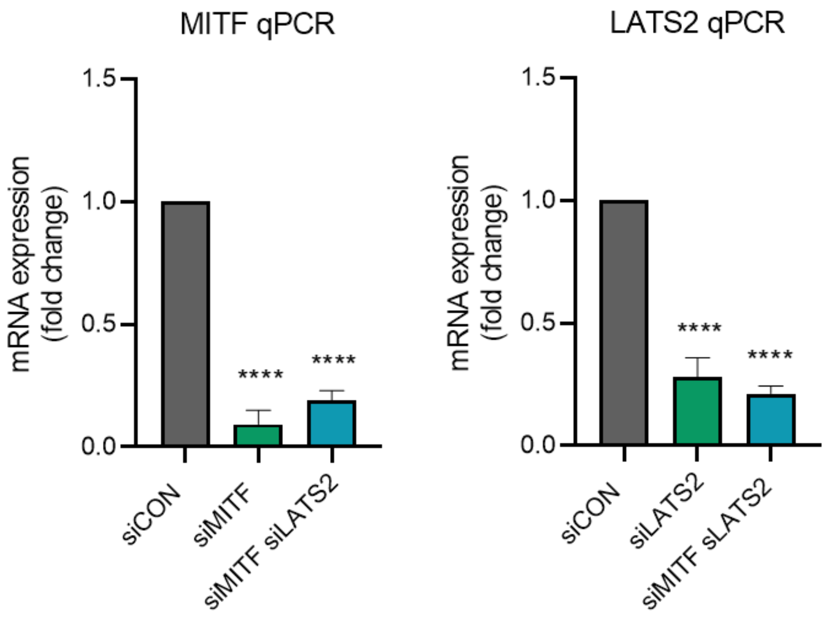


**E**

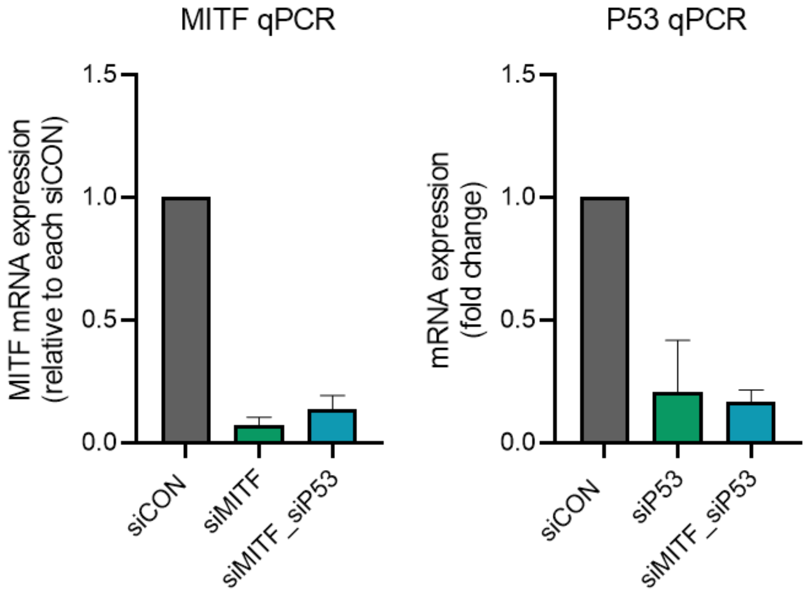


**Supplementary Figure S5 A** Cell cycle profiles of siCON, siMITF, siLATS2 and siMITF + siLATS2 treated U2OS cells. Cells were fixed after 48h siRNA treatment, followed by staining of DNA content with 7-aminoactinomycin D (7AAD) and flow cytometry analysis (unpaired t-test, n=4). **B** Confocal microscopy images (left) and quantification (right) of P53 protein levels in 624mel cells 48h after treatment with siCON, siMITF, siLATS2 and siMITF + siLATS2, cells were fixed and immunostained with a P53 specific antibody and DAPI to visualize nuclear cells. Scale: 20µM (one-way-ANOVA, n=3). **C** Cell cycle profiles of siCON and siMITF treated 624mel cells. Cells were fixed after 48h siRNA treatment, followed by staining of DNA content with 7-aminoactinomycin D (7AAD) and flow cytometry analysis (unpaired t-test, n=3). **D** MITF and LATS2 mRNA expression in indicated cell lines. Cells were treated with siRNA targeting siMITF, siLATS2, siMITF + siLATS2 or siRNA control for 48h, followed by real time qPCR analysis. (unpaired t-test, n=3). **E** MITF and P53 mRNA expression in indicated cell lines. Cells were treated with siRNA targeting siMITF, P53, siMITF + siP53 or siRNA control for 48h, followed by real time qPCR analysis. (unpaired t-test, n=3). Data presented as mean ± SD. **P* <0,05, ***P* <0,01, ****P* <0,001, *****P* <0,0001.

### Figure s6

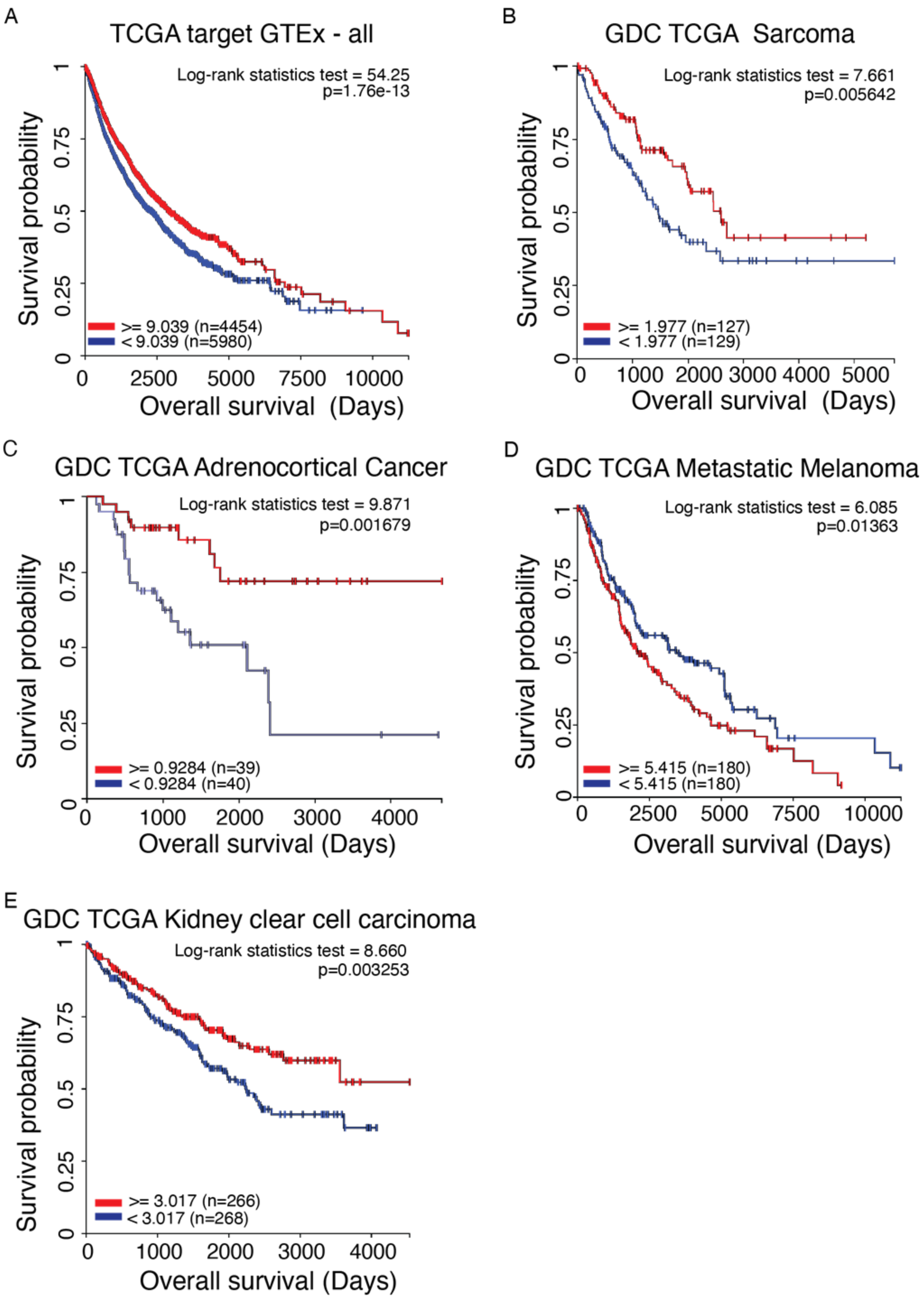
A
TCGA target GTEx - all
Log-rank statistics test = 54.25
p=1.76e-13
Survival probability
0
0.25
0.5
0.75
1
>= 9.039 (n=4454)
< 9.039 (n=5980)
0
2500
5000
7500
10000
Overall survival (Days)
B
GDC TCGA Sarcoma
Log-rank statistics test = 7.661
p=0.005642
Survival probability
0
0.25
0.5
0.75
1
>= 1.977 (n=127)
< 1.977 (n=129)
0
1000
2000
3000
4000
5000
Overall survival (Days)
C
GDC TCGA Adrenocortical Cancer
Log-rank statistics test = 9.871
p=0.001679
Survival probability
0
0.25
0.5
0.75
1
>= 0.9284 (n=39)
< 0.9284 (n=40)
0
1000
2000
3000
4000
Overall survival (Days)
D
GDC TCGA Metastatic Melanoma
Log-rank statistics test = 6.085
p=0.01363
Survival probability
0
0.25
0.5
0.75
1
>= 5.415 (n=180)
< 5.415 (n=180)
0
2500
5000
7500
10000
Overall survival (Days)
E
GDC TCGA Kidney clear cell carcinoma
Log-rank statistics test = 8.660
p=0.003253
Survival probability
0
0.25
0.5
0.75
1
>= 3.017 (n=266)
< 3.017 (n=268)
0
1000
2000
3000
4000
Overall survival (Days)

**Supplementary Figure S6** Kaplan-Meier survival data generated using xena browser. In each graph MITF high expressing tumors are represented in red and MITF low expressing tumors are represented in blue (log-rank test). Each graph represents survival data from different tumor types: **A** all tumor types, **B** sarcoma, **C** adrenocortical cancer, **D** metastatic melanoma and **E** kidney clear cell carcinoma. TCGA datasets, p-values, cut-off value between high and low RNA expression, and number of patients are shown on graphs. RSE norm_count was used for TCGA target GTEx samples and FPKM for the other samples. Kaplan-Meier survival analyses were generated using the UCSC Xena browser (https://xena.ucsc.edu/).
