## Supplementary data S1 S2 for "MITF maintains genome stability in non-melanocytic cell lineages and suppresses Hippo pathway signaling"

Sanger sequencing data of isoform specific PCR products presented in Figure 1F.

#### U2OS cells, A-isoform 1

AGTACGCTGA GAGCAGTTGG CCTGGTCTCG GGACTTGATT GATAAGCCTC CGATAACCTC 61

CTCCAGTATG ACATCACGCA TCTTGCTACG CCAACAACTC ATGGGTGAAC AGATGCTGGA 121

CCAGGATCGC AGGGAGCAGC AGCAGAAGCT GCAGGCGGCC CAATTCATGC AACAAAGAGT 181

GCCCGTGAGT CAAACACCAT CCATAAGCGT CAGTGTGGCC ACCACCCTTC CCTCTGCCTC 241

GCAGGGGCCG ATGGAAGTCC TGAAGGTGCA AACCCACCTC AAAAACCCCA CCAAGTACCA 301

CATACAGCAA GCCCAACGGC AGCAGGTAAA GCAGTACCTT TCTACCACTT TATCAAATAA 361

ACATGCCAAC CAAGTCCTGA GCTTGCCATG TCCAAACCAG CCTGGCGATC ATGTCATGCC 421

ACCGGTGCCG GTCAGCAGCG CACCCAACAG CCACATGGCT ATGCTTACGC TTAACTCCAA 481

CTGTGAAAAA GAGGGATTTT ATAAGTTTGA AGAGCAAAAC AGGGCAGAGA GCGAGTGCCC 541

AGGCATGAAC ACACATTCAC GAGCGTCCTG TATGCAGATG GATGATGTAA TCGATGACAT 601

CATTAGCCTA GAATCAAGTT ATAATGAGGA AATCTTGGGC TTGATGGATC CTGCTTTGCA 661

AATGGCAAAT ACGTTGCCTG TCTCGGGAAA TTGGATTTGA TCA //

#### SkMel28 cells, A-isoform 1

ATGTACGCTG AGAGCAGTTG CCTGTCTCGG GAACTTGATT GATCAGCCTC CGATAAGCTC 61

CTCCAGTATG ACATCACGCA TCTTGCTACG CCAGCAACTC ATGCGTGAGC AGATGCAGGA 121

GCAGGAGCGC ATGGAGCAGC AGCAGAAGCT GCAGGCGGCC CAGTTCATGC AACAGAGAGT 181

GCCCGTGAGT CAGACACCAG CCATAAACGT CAGTGTGCCC ACCACCCTTC CCTCTGCCAC 241

GCAGGTGCCG ATGGAAGTCC TTAAGGTGCA GACCCACCTC GAAAACCCCA CCAAGTACCA 301

CATACAGCAA GCCCAACGGC AGCAGGTAAA GCAGTACCTT TCTACCACTT TAGCAAATAA 361

ACATGCCAAC CAAGTCCTGA GCTTGCCATG TCCAAACCAG CCTGGCGATC ATGTCATGCC 421

ACCGGTGCCG GGGAGCAGCG CACCCAACAG CCCCATGGCT ATGCTTACGC TTAACTCCAA 481

CTGTGAAAAA GAGGGATTTT ATAAGTTTGA AGAGCAAAAC AGGGCAGAGA GCGAGTGCCC 541

AGGCATGAAC ACACATTCAC GAGCGTCCTG TATGCAGATG GATGATGTAA TCGATGACAT 601

CATTAGCCTA GAATCAAGTT ATAATGAGGA AATCTTGGGC TTGATGGATC CTGCTTTGCA 661

AATGGCAAAT ACGTTGCCTG TCTCGGGAAA CTTGATTGAT CA //

#### SkMel28 cells, M-isoform 1

TGGGAGGGAT AGTCTACCGT CTCTCACTGG GATTGGTGCC ACCTAAAACA TTGTTATGCT 61

GGAAATGCTA GAATATAATC ACTATCAGGT GCAGACCCAC CTCGAAAACC CCACCAAGTA 121

CCACATACAG CAAGCCCAAC GGCAGCAGGT AAAGCAGTAC CTTTCTACCA CTTTAGCAAA 181

TAAACATGCC AACCAAGTCC TGAGCTTGCC ATGTCCAAAC CAGCCTGGCG ATCATGTCAT 241

GCCACCGGTG CCGGGGAGCA GCGCACCCAA CAGCCCCATG GCTATGCTTA CGCTTAACTC 301

CAACTGTGAA AAAGAGGGAT TTTATAAGTT TGAAGAGCAA AACAGGCAGA GAGCGAGTGC 361

CCAGGCATGA ACACACATTC ACGAGCGTCC TGTATGCAGA TGGATGATGT AATCGATGAC 421

ATCATTAGCC TAGAATCAAG TTATAATGAG GAAATCTTGG GCTTGATGGA TCCTGCTTTG 481

CAAATGGCAA ATACGTTGCC TGTCTCGGGA ACTGCATTTG ATAAA //

### **Supplementary data S2**

qPCR primers used for qPCRs in the study.

| | |
|---|---|
| *MITF* | Forward _TTCCCACAGAGTCTGAAGCAAG_ |
| | Reverse _TCCAGCGCATGTCTGGATCA_ |
| *P53* | Forward _ CTTCCCTGGATTGGCAGC_ |
| | Reverse _ TTTCAGGAAGTAGTTTCCATAGGT_ |
| *LATS2* | Forward _CAGGATGCGACCAGGAGATG_ |
| | Reverse _AGGTCTGCTTAATGACCCGC_ |
| MITF - isoform A specific | Forward _TGAAGAGCCCAAAACCTATTACGA_ |
| | Reverse _GATCAATCAAGTTTCCCGAGACAG_ |
| MITF - isoform M specific | Forward _CCTTCTCTTTGCCAGTCCATCTTC_ |
| | Reverse _GATCAATCAAGTTTCCCGAGACAG_ |
| MITF - isoform C specific | Forward _CTTCAGTGGTTTTCCCACGAGCT_ |
| | Reverse _GATCAATCAAGTTTCCCGAGACAG_ |
| MITF - isoform H specific | Forward _GGAGGCGCTTAGAGTTCAGATG_ |
| | Reverse _GATCAATCAAGTTTCCCGAGACAG_ |
